# Calibration free counting of low molecular copy numbers in single DNA-PAINT localization clusters

**DOI:** 10.1101/2021.08.17.456678

**Authors:** J. Stein, F. Stehr, R. Jungmann, P. Schwille

## Abstract

Single-Molecule Localization Microscopy (SMLM) has revolutionized light microscopy by enabling optical resolutions down to a few nanometer. Yet, localization precisions commonly not suffice to visually resolve single subunits in molecular assemblies or multimeric complexes. Since each targeted molecule contributes localizations during image acquisition, molecular counting approaches to reveal the target copy numbers within localization clusters have been continuously proposed since the early days of SMLM, most of which rely on preliminary knowledge of the dye photo-physics or on a calibration to a reference. Previously, we developed localization-based Fluorescence Correlation Spectroscopy (lbFCS) as an absolute ensemble counting approach for the SMLM-variant DNA-Points Accumulation for Imaging in Nanoscale Topography (PAINT), for the first time circumventing the necessity for reference calibrations. Here, we present a revised framework termed lbFCS+ which allows absolute counting of copy numbers for individual localization clusters in a single DNA-PAINT image. In lbFCS+, absolute counting in individual clusters is achieved via precise measurement of the local hybridization rates of the fluorescently-labeled oligonucleotides (‘imagers’) employed in DNA-PAINT imaging. In proof-of-principle experiments on DNA origami nanostructures, we demonstrate the ability of lbFCS+ to truthfully determine molecular copy numbers and imager association and dissociation rates in well-separated localization clusters containing up to six docking strands. We show that lbFCS+ allows to resolve heterogeneous binding dynamics enabling the distinction of stochastically generated and *a priori* indistinguishable DNA assemblies. Beyond advancing quantitative DNA-PAINT imaging, we believe that lbFCS+ could find promising applications ranging from bio-sensing to DNA computing.

## 1 INTRODUCTION

The advent of super-resolution (SR) microscopy has revolutionized life science research by allowing to visualize specific biological structures at the nanoscale [1–4]. The SR methods summarized as SMLM, such as Photoactivated Localization Microscopy (PALM) [3], Stochastic Optical Reconstruction Microscopy (STORM) [4], and (DNA)-PAINT [5–7] circumvent the diffraction limit by acquiring image sequences of a ‘blinking’ target structure by stochastically activating only a small subset of all fluorescent labels at a time. Thus, these methods enable localization of individual dye molecules in each camera frame and downstream rendering of SR images from all obtained localizations. However, the limited photon budgets of dyes [8], imperfect labeling strategies and the physical size of the label (e.g. antibodies) cause these localizations to be scattered around the true position of the targeted molecule (forming a ‘localization cluster’) [9]. Within fixed cells, single molecules can thus only be pinpointed at lateral localization precisions of up to 10 nm [10], which is often not sufficient to reach molecular resolution and to visually resolve molecular complexes. To give an example, it is not possible to visually distinguish the two monomers within a dimer, because the localizations obtained from both molecules overlap within a localization cluster. However, since in SMLM each targeted molecule contributes a certain number of localizations to the SR image a quantitative analysis of the collected localizations from a specific (nanoscopic) volume in principle allows to infer back on the hidden number of targeted molecules within this volume [11].

Based on this concept, there has been a multitude of studies dedicated to the problem of ‘molecular counting’ since the early beginnings of SMLM, especially for the methods PALM and STORM [12–26]. While the required single molecule blinking in PALM is achieved by light-induced stochastic photoactivation and subsequent photobleaching of the fluorophores [3, 27], STORM exploits the light-induced photoswitching of fluorophores between a fluorescent bright state and a non-fluorescent dark state [4, 28]. Hence, for both methods, the success of a quantitative analysis of localization clusters critically depends on an exact photo-physical modeling of the specific system with respect to photobleaching [29], intrinsic and/or extended blinking [12, 14, 20] as well as photo-quenching [25] of the fluorophores in use.

In contrast to a direct and permanent dye labeling as used in STORM/PALM, DNA-PAINT exploits the transient hybridization of short single-stranded and fluorescently-labeled DNA probes (‘imagers’) to their complementary ‘docking strands’ attached as labels to the target molecules[6, 7]. Because the required blinking is generated by the stochastic imager-docking strand binding reaction, DNA-PAINT is largely independent of the photo-physical properties of fluorophores under appropriate experimental conditions (e.g. sufficiently low excitation intensities to reduce any residual photo-bleaching or the permanent photo-induced damage of docking strands) [7, 30–32]. In this case, localization clusters in DNA-PAINT data offer a unique potential for a quantitative interpretation, since the underlying bimolecular hybridization reaction between imager/docking strands is highly programmable and well-understood [11, 25, 33]. In fact, an approach termed quantitative PAINT (qPAINT) has been successfully used for molecular counting in localization clusters by using the imager influx rate as a calibration [34]. So far, all of the approaches to the problem of molecular counting in any of the SMLM variants were based on either (i) a priori knowledge of the blinking dynamics or the number of localizations per fluorescence marker (e.g., via supplementary experiments or theoretical modeling) or (ii) on an initial calibration directly within the sample by using isolated localization clusters originating from an assumed number of fluorescent molecules as a reference. Hence, those approaches only allow *relative* counting compared to a reference sample or given by the model assumptions.

In a previous study we introduced an approach termed lbFCS which allows *absolute* molecular counting in localization clusters in DNA-PAINT images, without the need of a separate reference measurement and using only minimal theoretical assumptions [32]. However, lbFCS required at least two measurements of the same sample at distinct and correctly adjusted imager concentrations, making an experiment rather tedious and time consuming. Additionally, lbFCS could only yield average values for both the underlying hybridization rates and the counted copy numbers and was hence not suited for the detection of possible heterogeneities between clusters. Finally, since its first implementation, several studies worked on the ‘speed-up’ of the DNA-PAINT reaction [10, 35–37] promising benefits on the achievable statistics (e.g. more binding events can now be recorded in the same amount of time).

Overcoming these limitations, we here present a revised framework lbFCS+ which allows extraction of *absolute* molecular numbers and hybridization rates of *single* DNA-PAINT clusters requiring only a *single* DNA-PAINT image acquisition. In proof-of-principle experiments on DNA origami nanostructures [38], we demonstrate the ability of lbFCS+ to truthfully determine molecular copy numbers and dissociation/association rates *k*_off_ & *k*_on_ of the imager/docking strand reaction in well-separated localization clusters containing up to six docking strands. We further thoroughly asses its applicable working range for reliable counting which is largely determined by the experimentally used imager concentrations and image acquisition length. Using lbFCS+ we could clearly give proof of changes in the imager/docking strand binding dynamics solely induced by placing docking strands at different positions of the DNA origami. Exploiting this effect we were even able to resolve heterogeneous binding dynamics within individual DNA-PAINT clusters allowing for the distinction of stochastically generated and a priori indistinguishable DNA assemblies.

## 2 MATERIALS AND METHODS

### 2.1 Brief recap of SMLM & DNA-PAINT binding dynamics

This section reviews the fundamental principles of DNA-PAINT binding kinetics, which constitute the basis of lbFCS+. For a detailed description of the working principles of SMLM in general and DNA-PAINT in particular, the reader is referred to [33] and [7], respectively.

A DNA-PAINT experiment [6, 7] is characterized by the transient binding reaction of short fluorescently-labeled DNA oligonucleotides in solution (‘imager strands’, short: ‘imagers’) to complementary ‘docking strands’ which are attached as labels to the target molecules of interest (see schematic in **Fig.1a**). At a given imager concentration *c* (typically on the order of ≈ 10 nM), the binding and unbinding reaction between imagers and docking strands is governed by the association rate *k*_on_ and the dissociation rate *k*_off_. While the dissociation reaction is a zero-order chemical reaction and thus independent of the reactant concentrations, the association reaction leading to the formation of the docking/imager strand duplex is a first order chemical reaction. Due to the ‘infinite reservoir’ of imagers in solution their concentration can be assumed to be constant during DNA-PAINT image acquisition. This leads to a constant effective association rate 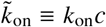 dependent on the imager concentration.

**Figure 1:**
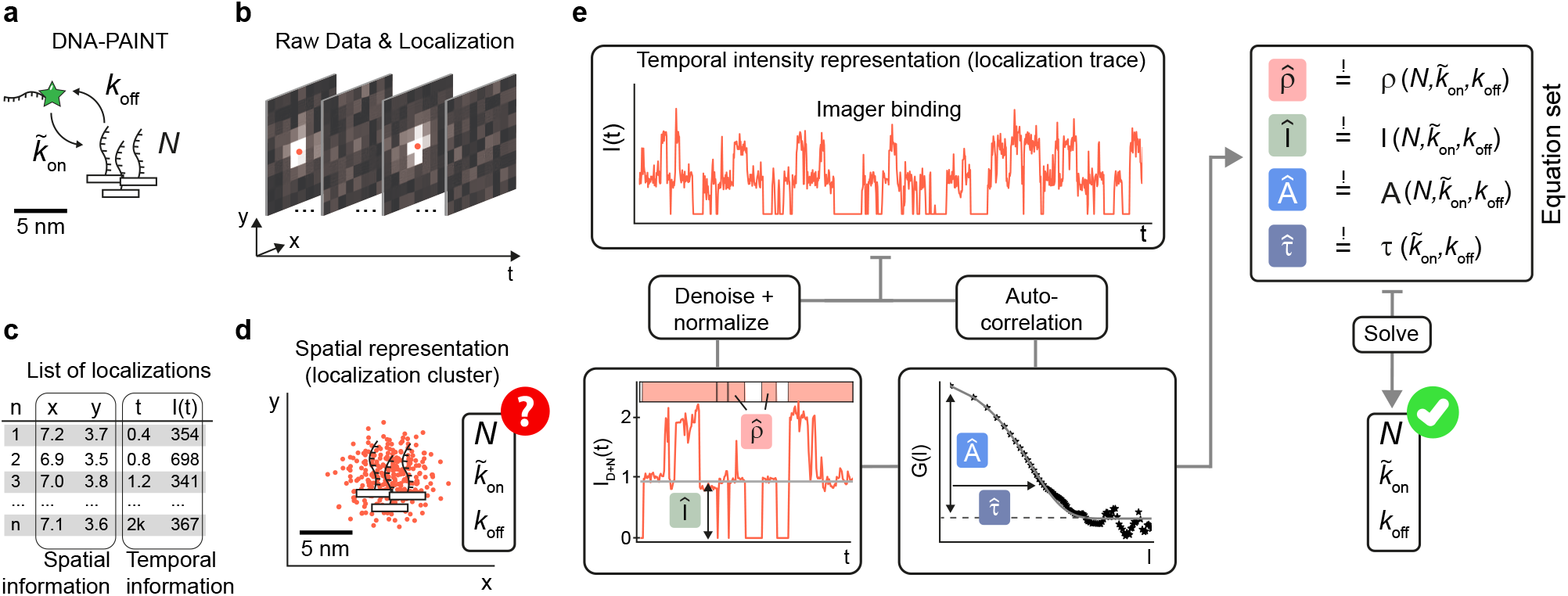
Absolute counting of molecular copy numbers in a single DNA-PAINT experiment. (**a**) Schematic of DNA-PAINT. The transient binding of fluorescently-labeled imager strands to complementary docking strands attached to the target molecules of interest. The binding reaction is governed by reaction rates *k*_off_ and 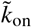. Multimeric organization of the target molecules can lead to accumulations of *N* docking strands at very close spacing (< 5 nm). (**b**) Sparse transient imager binding ensures the detection of single-molecule fluorescence signals during data acquisitions of typically several thousand frames. Fitting the center coordinates of each single-molecule detection during post-processing allows to obtain a localization that pinpoints the actual position of the underlying docking strand at nanometer precision (red points). (**c**) A processed DNA-PAINT data set consists of a list that contains all n obtained localizations (typically on the order of 10^6^). Each localization is associated to accessible quantities such as its spatial coordinates (x,y; in case of a 2D image), the frame *t* in which it was localized and its intensity *I*(*t*) as the number of detected photo-electrons. (**d**) An x-y scatter plot allows to render a super-resolved image as the spatial representation of the localization list (red dots). Ideally, the position of each docking strand is revealed by a clearly-distinguishable localization cluster. In case of multimeric targets, however, localizations obtained from multiple tightly-spaced docking strands can overlap in a not-resolvable localization cluster in the DNA-PAINT image. We are asking the question whether it is possible to derive the unknown physical quantities *N*, 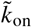 and *k*_off_ based on a single localization cluster, but for all localization clusters contained in the overall DNA-PAINT image. (**e**) Starting point of our solution to this problem is the intensity vs. time information that is associated to each localization cluster (compare (**c**)). This ‘intensity trace’ contains the temporal intensity fluctuations due to imager binding and unbinding that were detected from the position of the localization cluster during data acquisition. The intensity trace of each localization cluster is subject to two parallel analysis work streams (see 2.3 for detailed description): i) *Denoising* & *Normalization* which yields the two observables mean intensity *Î* and occupancy 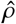 of the intensity trace as well as ii) *Autocorrelation* analysis which yields the two observables amplitude *Â* and decay constant 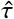 of the computed autocorrelation curve. Lastly, the four observables 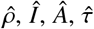 are input to a least square optimization of a defining set of equations to find a solution for the unknowns *N*, 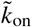 & *k*_off_

The schematic in **Fig.1a** depicts *N* docking strands spaced at only a few nanometer due to an exemplary local assembly of target molecules within the sample. During image acquisition, the ‘blinking’ raw signal recorded over time from this position consists of a series of bright frames (at least one imager strand bound to any of the *N* docking strands) and dark frames (no imager strand bound to any of the *N* docking strands), as illustrated in **Fig.1b**. As common for SMLM, the raw signal is converted into a list of localizations during post-processing, commonly referred to as super-resolution reconstruction. This is achieved by fitting a 2D Gaussian function to each of the identified diffraction limited spots, thereby pinpointing the fluorophore’s center coordinates (i.e. localization; red dots in **Fig.1b**). An exemplary list of localizations as obtained from the *N* docking strands by this procedure is shown in **Fig.1c**. Each localization carries information about the *x* and *y* coordinates of the identified spot (i.e. its spatial information) and a time stamp *t* of its occurrence as well as the total amount of recorded photo-electrons *I*(*t*) contained within the spot (i.e. its temporal intensity information). The spatial information contained in this list can be used to reconstruct a super-resolved image from our exemplary molecular assembly in form of a *x* – *y*-scatter plot for all localizations (see **Fig.1d**). However, due to the close docking strand spacing below the achievable localization precision it is not possible to visually distinguish individual docking strands and localizations overlap within the localization cluster.

### 2.2 Counting single molecules in DNA-PAINT localization clusters

The central question to which lbFCS+ aims to provide an answer is depicted in **Fig.1d**. In cases of molecular assemblies such as multimers, the spatial representation of localizations in a DNA-PAINT image often cannot reveal how many docking strands *N* are contained within a single localization cluster. Furthermore, the spatial representation does not reflect in any sense on the temporal information of imager binding (as given by 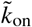 & *k*_off_) to the docking strands during image acquisition. Therefore, the quantities *N*, 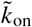 & *k*_off_ for each localization cluster must be considered unknown. It might further be the case that localization clusters feature distinct values in both *N* (e.g. due to varying degrees of multimerization or docking strand labeling efficiency) and in 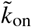 & *k*_off_ (e.g. due to the local sample environment affecting the imager accessibility). Hence, an ensemble measurement of 〈*N*〉, 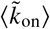 & 〈*k*_off_〉 would not be able to detect existing heterogeneities in either variable within the sample.

In contrast, the temporal intensity representation of a single localization cluster *I*(*t*) (referred to as ‘intensity trace’; see **Fig.1e**) contains the full information of imager binding under appropriately chosen experimental conditions during image acquisition. These conditions include that the reaction of imager-docking strand binding is at equilibrium (e.g. constant imager concentration *c* and temperature). Sufficiently low excitation intensities have to be employed in order to reduce possible photo-physical artifacts to a minimum such that the binding/blinking kinetics are solely determined by the hybridization rates 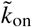 & *k*_off_. Explicitly, fluorophore photobleaching of bound imagers needs to be avoided by adjusting laser excitation with respect to *k*_off_ [32, 39]. Similarly, the photo induced loss of docking strands [30, 32] (leading to a decrease in *N* over the measurement time) has to be countered by appropriate measures (low excitation intensities, oxygen scavenger systems). Given that the stated conditions are fulfilled, lbFCS+ is able to find a separate solution *N*, 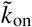 & *k*_off_ for individual localization clusters solely based on the information contained in their intensity traces.

We want to highlight that the applicability of lbFCS+ is intrinsically limited to targets which give rise to *distinct* and *well-separated* localization clusters in a DNA-PAINT image. Potential cellular targets are well-separated target molecule assemblies such as nuclear pore complexes in the nuclear envelope. We have previously published a python package, which allows automated detection and isolation of all localization clusters within a DNA-PAINT image [39] (available at https://doi.org/10.5281/zenodo.4792396), constituting the starting point of lbFCS+ analysis.

### 2.3 The analysis approach of lbFCS+

The concept of the lbFCS+ analysis framework is illustrated in **Fig.1e** and is applied in parallel to all detected localization clusters in a DNA-PAINT image. In case of a localization cluster originating from multiple docking strands, the intensity trace *I*(*t*) can show intensity fluctuations depending on the number of bound imagers at each time point. First, a step preserving non-linear de-noising filter [40] is applied to the intensity trace to generate a close to step-like behavior according to the number of simultaneously bound imagers (see **Supplementary Fig.1a**). Next, the de-noised intensity trace is normalized by the intensity recorded when only a single imager was bound (i.e. to the first intensity level). Hence, after normalization first level intensity values have a unit-less value of ‘1’ instead of an arbitrary photo-electron count, the second level has a value of ‘2’ and so forth. Further details about the normalization procedure are illustrated in **Supplementary Fig.1b**. Based on the de-noised and normalized intensity trace *I*_*D*+*N*_(*t*) the occupancy 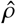 and the mean intensity 〈*I*_*D*+*N*_(*t*)〉 = *Î* are computed (see lower left panel in **Fig.1e**). The occupancy 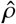 corresponds to the total time a signal was recorded at the position of the localization cluster divided by the total measurement time, i.e. the fraction the intensity trace was in a fluorescing state.

Analytic expressions for both 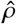 and *Î* can be derived under the assumption of equal and independent binding with 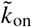 & *k*_off_ of each imager strand to each of the *N* docking strands. Then, the probability *P_k_* to find *k* imager strands simultaneously bound to *N* docking strands at an arbitrary point in time is given by a binomial distribution:

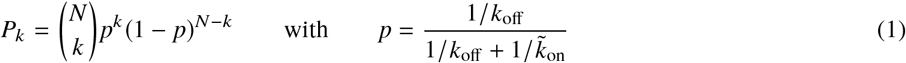

Intuitively, *p* corresponds to the probability to find a single docking strand in a fluorescing state, i.e. with an imager bound. The occupancy 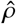 then corresponds to the inverse of the probability *P*_0_ of no imagers bound to the *N* docking strands:

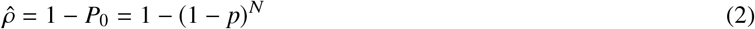

The mean intensity *Î* is simply given by:

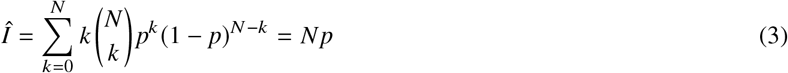

Hence, the expressions for both 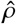 and *Î* solely depend on the unknowns *N*, 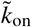 & *k*_off_.

Second, the autocorrelation function *G*(*l*) of the original localization trace *I*(*t*) is computed and *G*(*l*) is fitted with a mono exponential decay function yielding the amplitude *Â* and the characteristic decay time 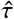 (see lower right panel in **Fig.1e**). This autocorrelation analysis step is analogous to our previous work lbFCS [32]. Again, both *Â* and 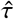 have known analytic expressions that solely depend on the unknowns *N*, 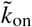 & *k*_off_ [32, 41]:

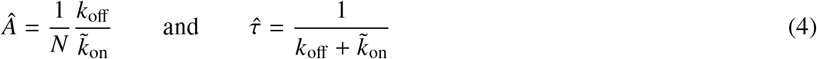

In the final step, the four observables 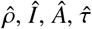 derived from the localization trace of each cluster are fed into the defining set of equations and a solution for the unknowns *N*, 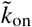 & *k*_off_ is found using least square optimization (see right panel in **Fig.1e**). We want to highlight that, using this approach we are only able to find a solution for the effective association rate 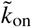 which is dependent on the imager concentration *c*. However, we can get an ‘concentration independent’ 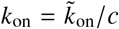 inserting the imager concentration *c* to which the sample was adjusted during sample preparation. Note that the thus derived ‘concentration independent’ *k*_on_ is still prone to pipetting errors, which can only be solved by independent concentration measurements or calibration to a reference sample (see Section 2.4).

We provide a `lbFCS+` python package (avaible at https://doi.org/10.5281/zenodo.5171076) that automatically computes the solutions for *N*, 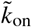 & *k*_off_ for all previously detected localization clusters in a DNA-PAINT image.

### 2.4 Materials and sample preparation

This study exclusively features DNA-PAINT experiments on DNA origami as synthetic targets mimicking molecular assemblies of docking strands. Rectangular DNA origami structures were designed using the ‘Design’ module of the Picasso software package [7]. Docking strand sequences were ‘5xCTC’ (CTCCTCCTCCTCCTC) and ‘Pm2*’ (TCCTC-CTC). Docking strand extended oligos were ordered from IDT. The imager of sequence ‘Pm2’ [32] (GAGGAGGA-Cy3b) was ordered from Eurofins. The adapter sequences ‘A20’ (AAGAAAGAAAAGAAGAAAAG) and ‘A20*+5xCTC’ (CTTTTCTTCTTTTCTTTCTT_TT_CTCCTCCTCCTCCTC) were ordered from IDT.

The folding reaction mix of each DNA origami design was prepared using 10× folding buffer (100 mM Tris,10 mM EDTA pH 8.0, 125 mM MgCl2; Ambion) and the following components: single-stranded M13 bacteriophage DNA scaffold p7249 (0.01 μM; Tilibit, M1-11), core staples (0.1 μM; ordered form Eurofins), biotin staples (0.01 μM; ordered form Eurofins), Docking strands (1 μM), 1× folding buffer in a total of 50 μl for each folding reaction. Annealing of was achieved via cooling the mixture from 80 to 25 °C in 3 h in a PCR thermocycler. A complete listing of the sequences of the cores staples and biotin staples for the rectangular DNA origami design can be found in the supplementary information of [7].

Standard DNA-PAINT reagents were ordered and prepared according to [7]: buffer A (10 mM Tris-HCl pH 7.5, 100 mM NaCl; Ambion), buffer B (5 mM Tris-HCl pH 8.0, 10 mM MgCl2, 1 mM EDTA; Ambion), BSA-biotin (Sigma-Aldrich, A8549) diluted @ 1 mg/ml in buffer A and streptavidin (Thermofisher, S-888) diluted @ 1 mg/ml in buffer A. 8-well microscopy slides (Ibidi, 80826) were plasma-cleaned for 1 min then washed 1× with 200 *μ*l buffer A. Each well could be used for an individual experiment, as explained in the following. 200 μl of BSA-biotin solution was flushed into the well, incubated for 2 min, removed and washed 1× with 200 *μ*l buffer A. Next, 200 μl of streptavidin solution was flushed into the well, incubated for 2 min, removed and washed 1× with 200 *μ*l buffer A and subsequently 1× with 200 *μ*l buffer B. DNA origami solution (diluted 1:200 in buffer B after folding) was flushed into the well, incubated for 5 min, removed and washed 2× with 200 *μ*l buffer B. Lastly, the desired imager strand concentration was directly adjusted in the well first adding the required amount of buffer B. As mentioned in Section 2.3, pipetting errors directly translate into the obtained result for *k*_on_. Therefore, all samples contained a subpopulation of reference origami consisting of *N* = 1 origami carrying a single Pm2* docking strand. Since Pm2* is a subset of our standard docking strand 5xCTC, the same imager Pm2 binds to both Pm2* (reference) and 5xCTC (target) docking strands but at a lower *k*_on_ (repetitive docking strands such as 5xCTC with multiple imager binding sites increase *k*_on_ [10, 37, 39]). Therefore, Pm2 reference localization clusters could be easily separated from 5xCTC clusters by using the occupancy 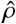 during analysis. After separation, resulting variations in *k*_on_ as obtained from Pm2 reference localization clusters were used as global calibration of the imager concentration.

### 2.5 Imaging

Imaging of DNA origami samples was performed on a custom-built flat-top TIRF microscope described in previous studies [31, 32, 39]. All fluorescence microscopy data was recorded with a sCMOS camera (2048 × 2048 pixels, pixel size: 6.5 μm; Andor Zyla 4.2). The camera was operated with the open source acquisition software *μ*Manager [42] at 2×2 binning and cropped to the center 700 × 700 pixel FOV. The exposure time was set to 400 ms corresponding to the aquisition duty cycle. The read out rate was 200 MHz and the dynamic range was set to 16 bit. The nearly homenous excitation irradiance [31] at the sample was set to ≈ 10 W/cm^2^. For detailed imaging parameters specific to the data presented in all main and supplementary figures refer to **Supplementary Table 1**.

## 3 RESULTS

### 3.1 Proof-of-principle demonstration of lbFCS+ on DNA origami

As in our previous work [32], we first tested lbFCS+ on DNA origami as synthetic targets that allow precise control of the number of docking strands per target. We designed four DNA origami variants carrying up to six docking strands *N* = 1, 2, 4, 6 of the sequence 5xCTC (≡ 5 repetitions of the triplet CTC). Note that here *N* is an upper bound due to the limited docking strand incorporation efficiency [43], i.e. a sample of *N* = 4 origami will also contain origami carrying only three, two or even one docking strand/s. We recorded a 30-min DNA-PAINT acquisition at 5 nM imager concentration for each origami variant immobilized on the cover glass of distinct wells of a microscopy slide (see Section 2.5 and **Supplementary Table 1** for detailed imaging conditions of all presented data). After localizing and rendering of the DNA-PAINT images, all localization clusters were automatically detected (see Section 2.3) and subject to lbFCS+ analysis.

**Fig.2a** shows the obtained counting results for *N* = 1 origami (# clusters = 1,994). The mean of the distribution at 〈*N*〉 = 1.06 is in close agreement with the expected value of 1, but indicates a slight tendency of over counting.

**Figure 2:**
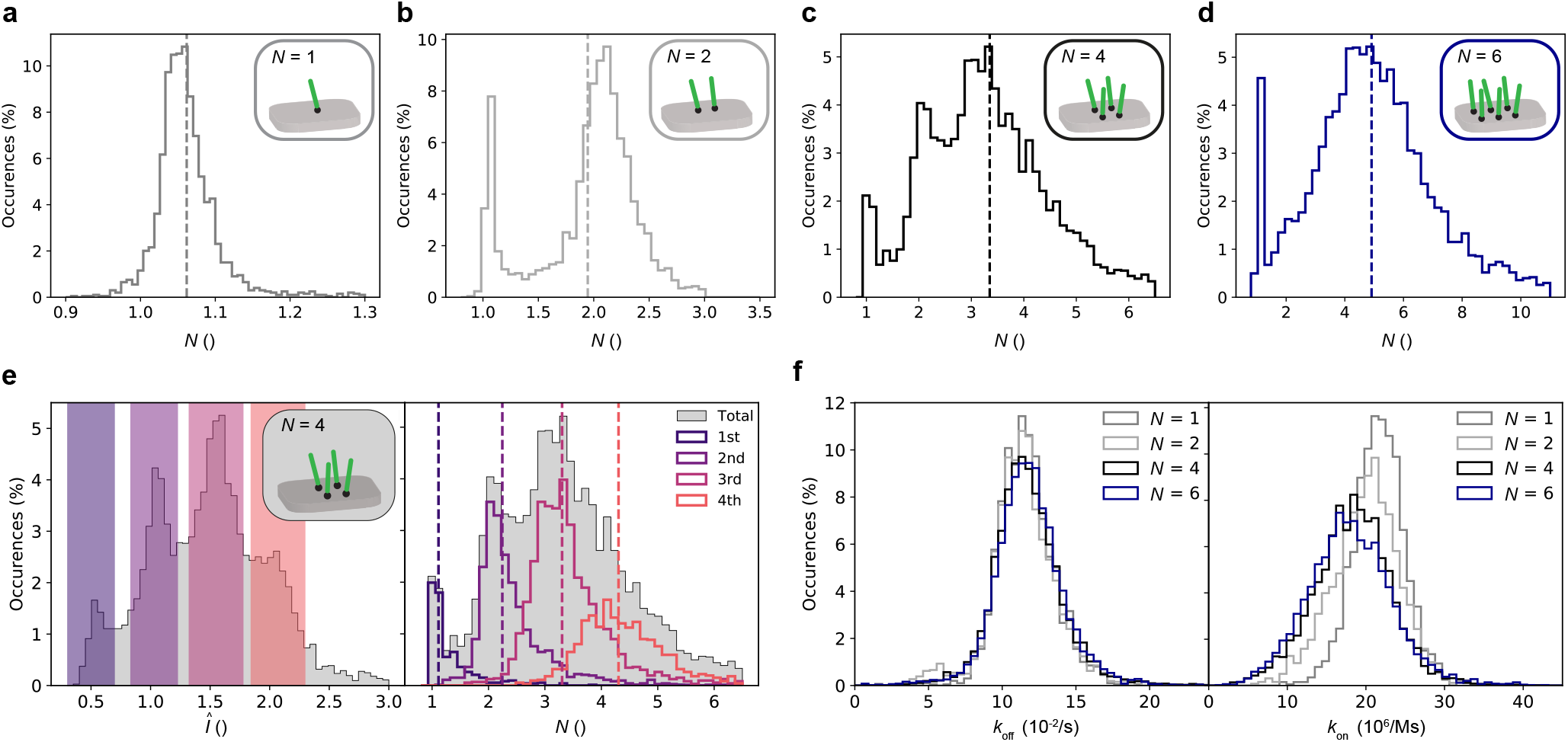
Proof-of-principle demonstration on DNA origami. (**a**) Counting results for DNA origami carrying a single docking strand (*N* = 1; # clusters = 1,994) with a mean of 〈*N*〉 = 1.06 (dashed line). (**b**) Same as (a) but for DNA origami carrying two docking strands (*N* = 2; # clusters = 2,582; 〈*N*〉 = 1.95). (**c**) Same as (a) but for DNA origami carrying four docking strands (*N* = 4; # clusters = 7,232; 〈*N*〉 = 3.35). (**d**) Same as (a) but for DNA origami carrying six docking strands (*N* = 6; # clusters = 5,038; 〈*N*〉 = 4.91). (**e**) *Î* distribution (grey; left panel) and *N* distribution (grey; right panel) for the *N* = 4 origami data set shown in (c). We defined subpopulations by selection of intervals in *Î* (colored intervals; left panel) and plotted their corresponding counting results *N* (colored solid lines) and mean values 〈*N*〉 (colored dashed lines, right panel). (**f**) Dissociation rates *k*_off_ (left panel) and association rates *k*_on_ (right panel) obtained via lbFCS+ analysis of all data sets shown in (a-d).

The counting results for the DNA-PAINT image of *N* = 2 origami (# clusters = 2,582) in **Fig.2b** features a prominent peak at *N* = 2 but also a smaller peak at *N* = 1 corresponding to origami with one of the two docking strands missing. 82 % of all localization clusters lie within 1.5 < *N* < 3 which corresponds to an average incorporation efficiency for any of the two docking strands of around 90 % (in good agreement with [43]).

The counting results obtained from the *N* = 4 origami image in **Fig.2c** (# clusters = 7,232) yielded a distribution with clearly distinguishable peaks located at *N* = 1, 2 and 3. Based on the mean of the distribution 〈*N*〉 = 3.35 (dashed line) we estimated a slightly lower incorporation efficiency of around 84 % [43]. However, we observed a broadening of the distribution toward higher *N*, hindering a visual distinction of the peak at *N* = 4.

This is further confirmed when looking at the counting results derived from the *N* = 6 origami data set (see **Fig.2d**). While for *N* >= 5 it is not possible to visually distinguish incremental copy numbers, the mean of the distribution at 〈*N*〉 = 4.91 (dashed line) still yields a reasonable ensemble average result (corresponding to an incorporation efficiency of around 82 %) [43].

Next, we turned back to the *N* = 4 data set in order to find out whether it is possible to achieve a clear distinction between *N* = 3 and *N* = 4. Remarkably, we found that the distribution of mean intensities *Î* obtained from all localization clusters exhibited four peaks, as depicted in the left panel of **Fig.2e**. Intuitively, the leftmost peak, i.e. the lowest mean intensity should correspond to *N* = 1 origami, since increasing numbers of docking strands lead to higher values of *Î* due to the increasing probability of simultaneous binding of multiple imagers (see Eq. 3). We confirmed this by selecting localization clusters lying within the colored intervals in *Î* and by comparing the corresponding subpopulations in *N* to the overall obtained distribution (**Fig.2e**, left and right, respectively). This selection in *Î* allowed us to obtain mean counting results for the subpopulations that are close to the expected values of *N* = 1, 2, 3 and 4 (colored dashed lines). The visual inspection of exemplary intensity traces from the selected intervals in *Î* confirms the applicability of this approach (see **Supplementary Fig.2**). Again, we observed a slight over counting that is more prominent for increasing *N*.

After inspection of the counting results, we turned our attention to the imager hybridization rates obtained via lbFCS+ analysis of the same four data sets as in **Fig.2a-d**. **Fig.2f** shows the corresponding *k*_off_ distributions (left) and *k*_on_ distributions (note that lbFCS+ yields 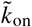, from which 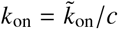 was calculated using the absolute imager concentration, see Section 2.1). Overall, *k*_off_ shows very good agreement for all four data sets independent of the number of docking strands per origami with a relative width of STD(*k*_off_)/〈*k*_off_〉 ≈ 18 %.

In *k*_on_, however, we observed broader distributions compared to *k*_off_ with relative widths STD(*k*_on_)/〈*k*_on_〉 increasing from 14 % for *N* = 1 to 30 % for N=6. Additionally, we observed a slight, but systematic decrease with increasing numbers of docking strands with 〈*k*_on_〉 decreasing from 22×10^6^ 1/Ms for *N* = 1 to 18×10^6^ 1/Ms for *N* = 6. As described in Section 2.4, reference origami allowed for a calibration of the imager concentrations to minimize pipetting errors affecting *k*_on_ (see **Supplementary Fig.3** for the calibration results of the *N* = 1, 2, 4, 6 datasets shown in **Fig.2a-d**).

Both the experimentally observed upper limit in distinguishing copy numbers of *N* >= 4 and the apparent dependence of *k*_on_ on *N* led us to further assess the applicable working range of lbFCS+.

### 3.2 Assessing the working range of lbFCS+

To investigate the applicable working range of lbFCS+ and possible systematic artifacts it was obligatory to perform the analysis on a data set of known ground truth. Therefore, we computationally combined arbitrary localization clusters from the *N* = 1 data set (see **Fig.2a**) into clusters of user-defined *N*_in_ = *k* × (*N* = 1), a concept we already applied in an earlier study [32]. This ‘regrouping’ is equivalent to the computational addition of intensity traces *I*(*t*) of the individual clusters. To exclude varying intensity levels between the original clusters (e.g., caused by speckles in the illumination profile) the intensity traces *I*(*t*) where normalized prior to addition (see **Supplementary Fig.1**). We want to highlight, that this procedure completely preserved the experimental intensity noise distribution.

**Fig.3a** shows the counting results obtained from computationally regrouped clusters consisting of up to eight experimental *N* = 1 clusters (i.e. *N*_in_ = *k* × (*N* = 1) up to *k* = 8). Overall, the counting results *N* agree very well with the expected *N*_in_. Again, we observed a slight systematic offset of the resulting means 〈*N*〉(colored dashed lines) towards higher *N*. Although this effect seemed to increase with *N*_in_ in absolute terms, the relative offsets ((〈*N*〉 – *N*_in_)/*N*_in_ remained constant at ≈ 6 % (compare to **Fig.2a**). Similar to the experimental results the width of the *N* distributions broadened with increasing *N*_in_ (compare to **Fig.2a-e**). The relative width of each distribution defined as STD(*N*)/〈*N*〉 increased proportional to 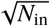, starting with a value of 4 % for *N* = 1. This broadening behavior is in line with the experimental *N* results displayed in **Fig.2a-d**. However, from our regrouping analysis we would expect that it should be possible to clearly distinguish the peaks between *N* = 3 and *N* = 4, which is not the case for DNA origami (see **Fig.2c**). This indicates an additional source of uncertainty of lbFCS+ counting towards higher *N*.

**Figure 3:**
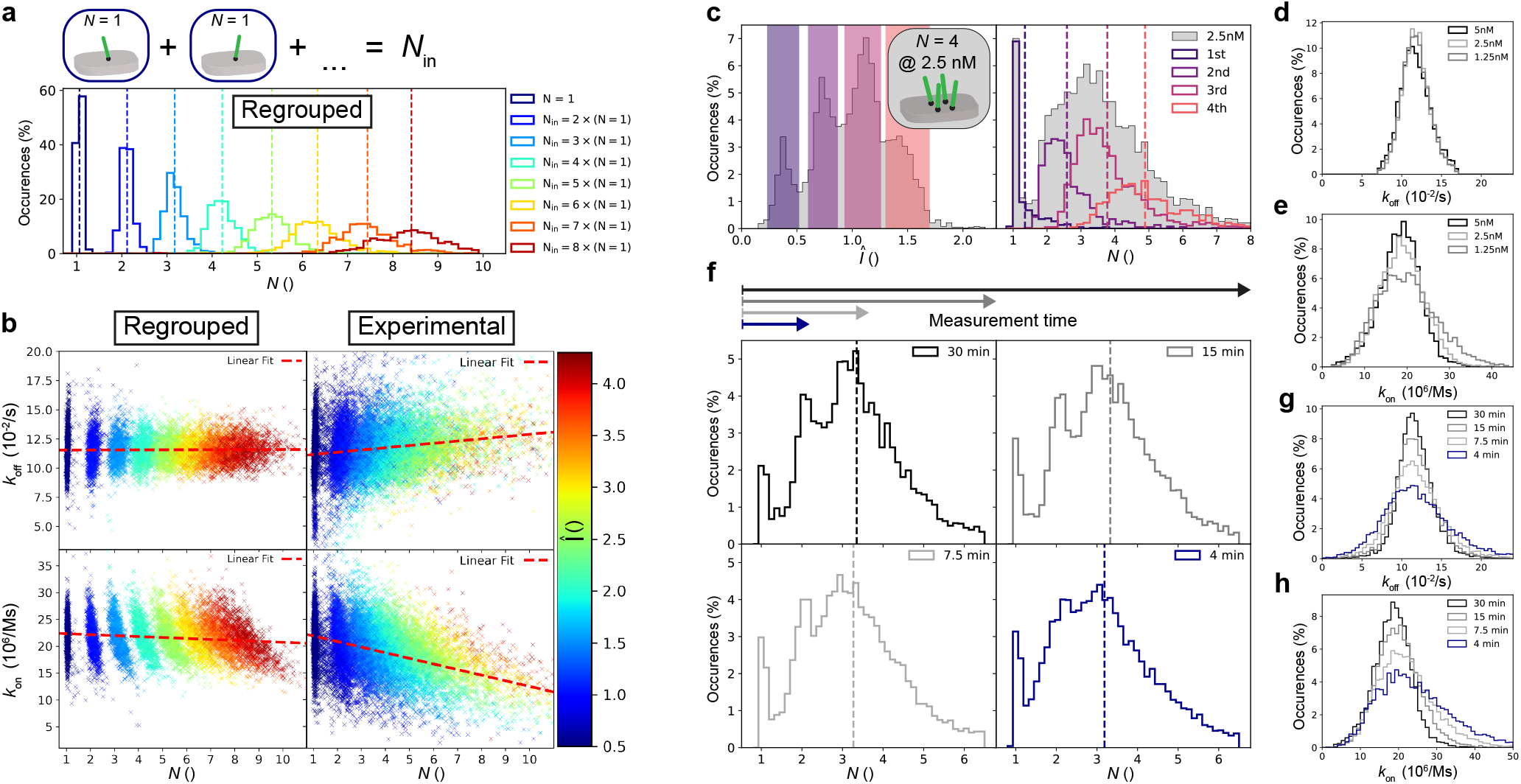
Assessing the working range of lbFCS+. Localization clusters from the *N* = 1 data set (see **Fig.2a**) were selected at random and subsequently computationally combined into clusters of user-defined *N*_in_ = *k* × (*N* = 1) and subject to lbFCS+ analysis (# clusters = 1,000 for each *N*_in_). **(a)** Counting results *N* obtained from computationally-combined clusters consisting of up to eight experimental *N* = 1 clusters (i.e. *N*_in_ = *k* × (*N* = 1) up to *k* = 8) and their corresponding means 〈*N*〉 (dashed lines). **(b)** Scatter plot *k*_off_ vs. *N* (upper panels) and scatter plot *k*_on_ vs. *N* (lower panels) for all computationally-combined clusters (left panels) shown in (a) and for all experimental clusters from *N* = 1, 2, 4, 6 origami (right panels) shown in **Fig.2a-d**. Each cluster was color-coded according to its mean intensity *Î* (which is linearly proportional to *N*, see Eq.3). The red dashed lines indicate the optimum linear fits over all data points in each panel. **(c)** Analysis analogous to **Fig.2e** for DNA origami containing four docking strands (*N* = 4,) but measured at a reduced imager concentration of 2.5 nM (# clusters =3,697). **(d)** Dissociation rates *k*_off_ obtained via lbFCS+ analysis for DNA origami containing four docking strands (*N* = 4) measured at varying imager concentrations (# clusters = 5,834 for 1.25 nM). **(e)** Association rates *k*_on_ for the same data sets as in (d). **(f)** Counting results for the *N* = 4 origami data set shown in **Fig.2c** for varying measurement times. The original data set (30 min measurement time) was reduced to the first 15 min, 7.5 min and 4 min, respectively, prior to analysis. The dashed lines indicate the mean 〈*N*〉 of each distribution. **(g)** Dissociation rates *k*_off_ obtained via lbFCS+ analysis for the data shown in (g). **(h)** Association rates *k*_off_ obtained via lbFCS+ analysis for the data shown in (g).

Next, we focused on the hybridization rates obtained from the regrouped data sets. The upper left panel of **Fig.3b** shows a scatter plot of the obtained *k*_off_ vs. *N* result for all localization clusters from the eight regrouped data sets. Each cluster was color coded using its respective mean intensity value *Î* (which is linearly proportional to *N*, see Eq.3). Contrarily to the broadening in *N*, we observed a narrowing of the *k*_off_ distributions for increasing *N*_in_. Both the shape of the distributions with respect to *k*_off_ and *N* as well as the negligible slope of a linear fit of all data points (red dashed line) indicated that the solutions for *k*_off_ and *N* are largely decoupled.

Similarly, the lower left panel of **Fig.3b** shows the analogous scatter plot for *k*_on_ vs. *N*. Here, the linear fit over all clusters exhibited a minor, but negligible decrease in *k*_on_ for increasing *N* with respect to experimental measurement errors (see description of **Fig.2f** in Section 3.1). Furthermore, we observed a hyperbolic shape of the *k*_on_ vs. *N* distributions for increasing *N*_in_. This is due to the fact that three of the four observables (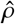, *Î* and *Â*) used as input to the final set of equations (see **Fig.1e**, right) are in first order approximation proportional/indirect proportional to the product 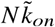 (see Eq.2,3 and 4). Additionally, the observable 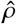 only contains valuable information for fluorescence traces featuring interruptions, thereby constituting an upper operational limit of lbFCS+ at a given imager concentration. For increasing *N*, almost uninterrupted intensity traces lead to saturation of 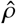 (e.g. 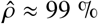 for *N*_in_ = 8) and as such to a loss of information for the defining set of equations. For this reason, lbFCS+ is designed for the application to targets containing low copy numbers of docking strands 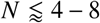 (depending on the used imager concentration and *k*_on_). To experimentally asses its upper limit we imaged and analyzed origami containing 12 docking strands, still yielding reasonable ensemble results when imaged at a concentration of 2.5 nM (see **Supplementary Fig.4**)).

To compare the findings obtained from computational regrouping to the experimental data we depicted the combined results from *N* = 1, 2, 4, 6 origami (see **Fig.2a-d**) in an analogous *k*_off_ vs. *N* scatter plot in the upper right panel of **Fig.3b**. In contrast to the computationally combined clusters, linear fitting over all experimental clusters (red dashed line) indicated an increase in *k*_off_ with *N*. On the other hand, the experimentally obtained *k*_on_ vs. *N* scatter plot in the lower right panel of **Fig.3b** similarly yields increasingly hyperbolic distributions as observed for the regrouped clusters. Linear fitting of all data points (red dashed line) showed a clear decrease in *k*_on_ with increasing *N*, as already observed in **Fig.2f**. This decrease in *k*_on_ with increasing *N* is significantly larger compared to the regrouped data sets (see lower left panel).

The observations in both *k*_on_ and *k*_off_ indicated that intensity traces recorded from origami containing multiple docking strands are not exactly equal to the simple addition of the individual single docking strand signals. We suspected that the docking strand position on the DNA origami could lead to local changes in *k*_on_ and *k*_off_, thereby possibly giving rise to a measurement bias for higher *N*. Remarkably, when performing four control experiments, each time only with one of the four docking strands of the *N* = 4 origami incorporated, we could not observe any position dependence neither in *k*_on_ nor in *k*_off_ (see **Supplementary Fig.5**). We hypothesize that cooperative binding due to the spatial proximity of the docking strands could be a possible explanation for this behavior. However, over the applicable range of 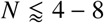, this does not affect the ability of lbFCS+ to obtain correct counting results and only causes minor deviations in the measured hybridization rates (≤ 2 % and ≤ 20 % in *k*_off_ and *k*_on_, respectively).

As a next step we assessed the lower limit of the applicable working range by measuring *N* = 4 origami at reduced imager concentrations. Analogous to **Fig.2e** it was still possible to clearly distinguish subpopulations in *Î* in order to identify the corresponding result in *N*, but at a reduced imager concentration of 2.5 nM (see **Fig.3c**). It was even possible to repeat the same analysis for a sample imaged at an imager concentration of 1.25 nM (see **Supplementary Fig.6**). The shapes of the *N* distributions are in close agreement with the results obtained from the computational combination of experimental *N* = 1 clusters measured at a concentration of 1.25 nM to clusters of defined *N*_in_ = *k* × (*N* = 1) as presented in **Supplementary Fig.7** (analogous to **Fig.3a,b**).

Surprisingly, imaging *N* = 4 origami at lower imager concentrations had no significant effect on the resulting *k*_off_ values as shown in **Fig.3d**. However, we observed a broadening in *k*_on_ with decreasing imager concentrations as expected from the broadening in *N* (see **Fig.3e**).

Finally, we investigated the effects of the measurement time on lbFCS+ analysis (i.e. the image aquisition time). We therefore reduced the original *N* = 4 data set (30 min measurement time) to the first 15 min, 7.5 min and 4 min prior to analysis. Remarkably, it was only possible to observe significant changes in the resulting *N* distribution at measurement times 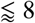 min (see **Fig.3f**). A measurement time of 4 min would correspond to an expectation value of only ≈ 13 imager binding events per single docking strand. We observed a broadening in both *k*_off_ and *k*_on_ for reduced measurement times as apparent from **Fig.3g** and **Fig.3h**, respectively. Finally, our optimization of the required measurement times allowed us to image 18 FOVs containing a total of ≈ 50.000 origami in ≈ 3 h of total measurement time, still yielding robust quantitative results (see **Supplementary Fig.8**).

### 3.3 Distinction of nanoscopic DNA assemblies via binding dynamics

Driven by the high accuracy of lbFCS+ to determine hybridization rates we hypothesized that it might be possible to distinguish DNA constructs via detection of slight changes in the position dependent imager-docking strand binding dynamics.

Besides the direct incorporation of 5xCTC docking strands as used in the preceding experiments, we designed DNA origami carrying a single 20 base ‘adapter’ docking strand of sequence A20 (see Section 2.4 for exact sequences). Addition of oligos carrying both the complementary adapter region and the docking strand sequence (A20*+5xCTC, referred to as ‘linker strand’) allowed us to permanently install the docking strand further away from the origami surface via the double-stranded link A20+A20* (≈ 10 nm at full elongation), as depicted in **Fig.4a**. We distinguished between the ‘Direct’ configuration (5xCTC incorporated; grey box) and the ‘Link’ configuration (5xCTC on top of the double-stranded A20 linker; orange box). We want to highlight that both configurations are not rigid but experience rotational freedom introduced by single stranded TT-spacers (black dots; one for Direct and two for Link, respectively).

**Figure 4:**
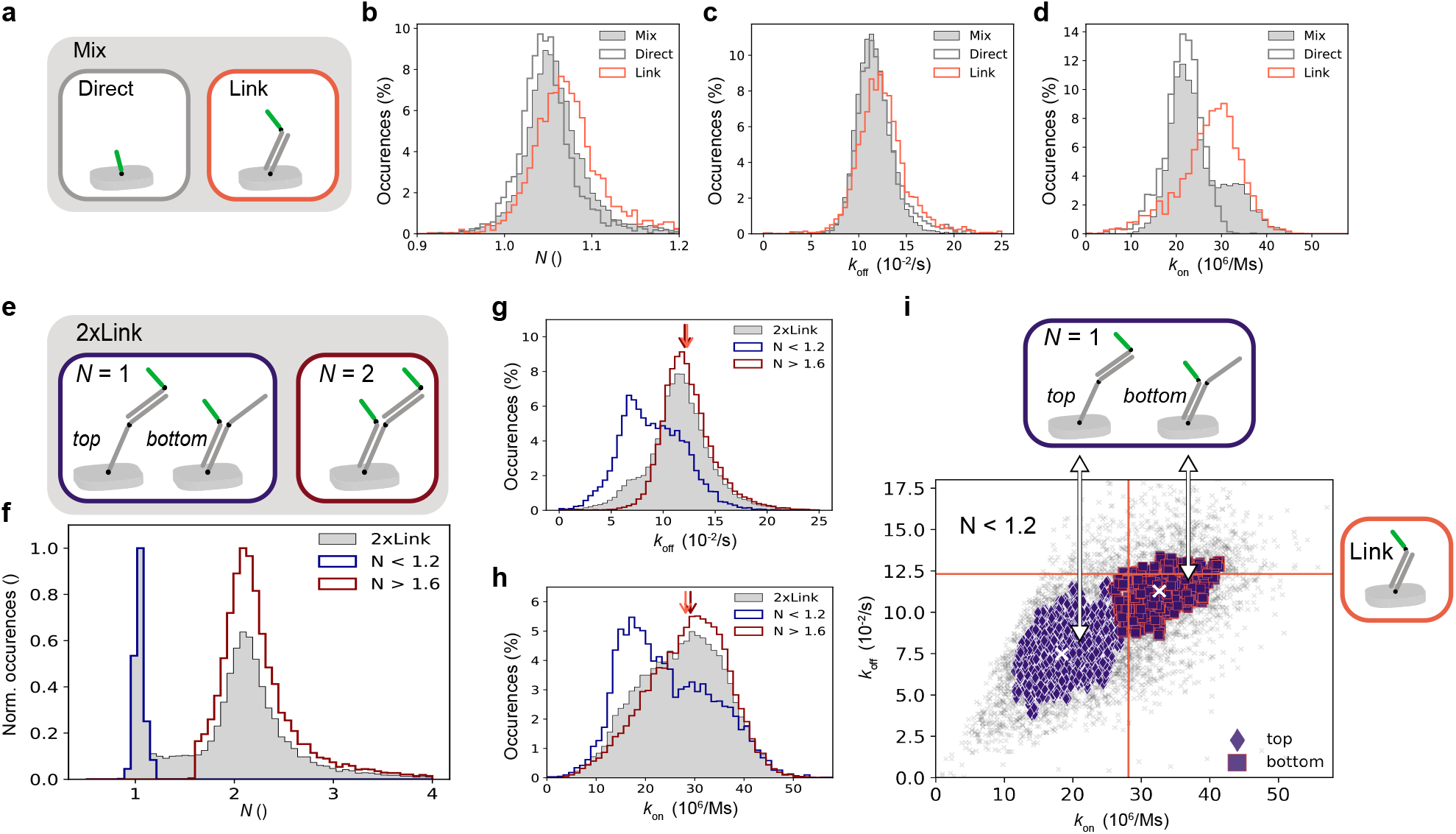
Distinction of nanoscopic DNA assemblies via binding dynamics. **(a)** Schematic of the direct incorporation of 5xCTC docking strands into DNA origami (‘Direct’; dark-grey box) and permanent attachment of 5xCTC docking strands via a ‘linker strand’ (A20*+5xCTC) after folding (‘Link’; orange box). Black dots indicate single stranded TT-spacers. We imaged three samples containing 1) only Direct origami, 2) only Link origami or 3) a mixture of both configurations (‘Mix’; filled grey box). **(b)** lbFCS+ counting results as obtained from the Direct origami, the Link origami and the mixed sample as illustrated in (a). Direct sample contained # clusters = 2,088; Link: # clusters = 2,780; Mix: # clusters = 9,136. **(c)** Dissociation rate *k*_off_ as obtained from the same data as shown in (b). **(d)** Association rate *k*_on_ as obtained from the same data as shown in (b). **(e)** DNA origami design featuring 2x linker binding sites (referred to as ‘2xLink’). After incubation with linker strands, 2xLink origami can be in three possible states (schematically depicted). The first two states consist of a single linker strand (i.e., *N* = 1) bound to the 2xLink origami at either the bottom or the top position (blue box). The third state corresponds to both A20 sites being occupied by a linker strand (i.e., *N* = 2; dark-red box). **(f)** Total *N* distribution obtained from imaging 4 distinct samples of 2xLink origami, which were incubated for 3 min at either 100 nM, 40 nM, 5 nM or 2 nM linker strand concentrations (**Supplementary Fig.9** shows separate results for each sample). For further analysis, we split the total distribution into into clusters yielding either *N* < 1.2 (blue) or *N* > 1.6 (dark red). Total # of 2xLink origami localization clusters from 4 data sets = 27,348; *N* < 1.2: # clusters = 6159; *N* > 1.6: # clusters =19,192. **(g)** Dissociation rate *k*_off_ as obtained from the same data as shown in (f). The dark red arrow indicates the median of the *N* > 1.6 subpopulation, the orange arrow indicates the median of the Link origami shown in (c). **(h)** Results analogous to (g) but for the association rate *k*_on_. **(i)** Scatter plot of *k*_off_ vs. *k*_on_ for all 2xLink origami yielding *N* < 1.2, see (f). Hierarchical density based clustering (hdbscan) [44] (used parameters: `metric = ‘12’, min_cluster_size = 500, min_samples = 8`) of the data yielded two groups. The median of each group is marked by a white cross and the corresponding median as obtained from Link origami shown in (c,d) is indicated by the orange lines. We assigned the bottom *N* = 1 state of the 2xLink origami to the upper right group (dark-blue squares) due to 1) the proximity of its median to the median of the Link origami and 2) its resemblance in design to the Link origami (compare orange box in (a) with blue box in (e)) Vice versa, we assigned the top *N* = 1 state of the 2xLink origami to the lower left group (dark-blue diamonds). All *N* < 1.2 origami contained # clusters = 6159; top: # clusters = 2630; bottom: # clusters = 981.

We subsequently imaged three samples containing 1) only Direct origami, 2) only Link origami or 3) a mixture of both configurations. lbFCS+ analysis yielded the expected number of docking strands *N* = 1 (see **Fig.4b**) in all cases and we could not observe any alteration of *k*_off_ between the different configurations (see **Fig.4c**). In contrast, we observed an increased *k*_on_ for the Link configuration compared to the Direct configuration (see **Fig.4d**). We suppose that both the increased mobility of the docking strand and larger distance from the origami surface promote a higher chance of imager binding for the Link configuration. This shift was large enough to clearly identify the Link/Direct origami in the bimodal *k*_on_ distribution of the sample containing both configurations (Mix).

Following the same reasoning, we designed DNA origami similar to the Link assembly (see **Fig.4a**) but now providing a second possible binding site for the A20 adapter (referred to as ‘2xLink’, see **Fig.4e**). Due to the stochastic nature of linker strand binding, this origami configuration can be observed in one of three possible states. The first two states consist of a single linker strand (i.e., *N* = 1) bound to the 2xLink origami at either the bottom or the top (blue box in **Fig.4e**). The third state corresponds to both A20 sites (i.e., *N* = 2) being occupied by a linker strand (dark-red box in **Fig.4e**). Since the ratio of origami in a *N* = 1 or *N* = 2 configuration should be manipulable via variation of linker strand concentrations, we imaged four samples of 2xLink origami which were previously incubated for 3 min at 100 nM, 40 nM, 5 nM and 2 nM linker strand concentrations.

**Fig.4f** shows the total *N* distribution obtained from lbFCS+ analysis of the four data sets (grey), confirming the expected counting result of either *N* = 1 or *N* = 2. While for a 100 nM linker strand concentration ≈ 80 % of all origami had bound two linkers, at 2 nM it was only ≈ 60 %, validating the concentration dependence during incubation on the probability of 2xLink origami to be found in an *N* = 1 or *N* = 2 state (see **Supplementary Fig.9**).

Again, we were interested in potential variations in the measured hybridization rates depending on the state of each 2xLink origami. For this reason, we isolated the *N* = 1 and *N* = 2 configurations by separating localization clusters that yielded either *N* < 1.2 (blue) or *N* > 1.6 (dark red), respectively (see **Fig.4f**). Already the total *k*_off_ distribution of all 2xLink origami (grey) presented in **Fig.4g** revealed two subpopulations located at *k*_off_ ≈ 7×10^-2^ 1/s and *k*_off_ ≈ 11×10^-2^ 1/s. These two subpopulations became especially prominent when looking at only *N* = 1 localization clusters (blue), confirming the existence of a top and a bottom state of *N* = 1 origami that give rise to a distinct *k*_off_. In contrast, the *N* = 2 clusters yielded a homogeneous *k*_off_ distribution with a median (dark red arrow) nearly identical to the median of previous Link origami (orange arrow, obtained from orange distribution shown in **Fig.4c**).

Inspection of the *k*_on_ results yielded a similar behavior, as depicted **Fig.4h**. While the total *k*_on_ distribution showed a somewhat broadened shape (grey), selection of *N* = 1 origami clearly revealed two subpopulations located at *k*_on_ ≈ 17×10^6^ M/s and *k*_on_ ≈ 31×10^6^ M/s. This suggested that for *N* = 1 origami the top and bottom states also give rise to a different *k*_on_. The *k*_on_ values obtained from *N* = 2 origami resulted in a broad distribution with a skew toward lower *k*_on_ values. However, its median (dark red arrow) was again close to the median in *k_on_* as obtained for the Link origami (orange arrow, obtained from orange distribution shown in **Fig.4d**).

In conclusion, this suggests that the signal from *N* = 2 origami is actually a superposition of heterogeneous signals due to the distinct binding dynamics of the top/bottom *N* = 1 states. Strictly speaking, here *N* = 2 origami hence violate our assumption of equal and independent binding rates used in the derivation of Section 2.3. The coincidence of the *N* = 2 peak (dark red) with the right peaks of the two possible *N* = 1 configurations (blue) indicates that in case of heterogeneous rates lbFCS+ analysis is dominated by the larger value in both *k*_off_ and *k*_on_ (see **Fig.4g** and **h**, respectively). Regardless of heterogeneous rates, lbFCS+ analysis still yielded the correct counting results (compare **Fig.4f**).

Finally, we aimed to exploit the heterogeneous binding kinetics to identify the (otherwise indistinguishable) top/bottom states within the *N* = 1 subpopulation. Indeed, hierarchical density based clustering (hdbscan) [44] allowed us to classify two distinct states in the scatter plot of *k*_off_ vs. *k*_on_ over all *N* = 1 localization clusters (see **Fig.4i**). Intuitively, the bottom *N* = 1 state of the 2xLink origami should be close to the Link configuration depicted in **Fig.4a**. Comparison of the median *k*_off_ & *k*_on_ of each class (white crosses) with the median *k*_off_ & *k*_on_ as obtained for the Link origami (orange lines, obtained from **Fig.4b** and **d**) hence allowed to associate the top right class (square) with the bottom *N* = 1 state and the top left class with the top *N* = 1 state (diamonds). We classified almost three times as many origami in a top state (# clusters = 2630) as in a bottom state (# clusters = 981) suggesting a lower binding probability of linker strands to the bottom position. Interestingly, we observed a significantly lower *k*_off_ (i.e. a longer binding duration) and a lower *k*_on_ for docking strands placed at the top position when compared to the bottom position.

## 4 DISCUSSION

In summary, lbFCS+ is, to our knowledge, the first method capable of extracting both absolute molecular copy numbers and DNA hybridization rates of individual DNA-PAINT localization clusters within a single DNA-PAINT image. Based on only minimal experimental requirements and theoretical assumptions it thus provides a solution to the long prevailing problem of ‘molecular counting’ in SMLM without the need of any initial calibration or modeling [11, 25, 33].

In proof-of-principle experiments on DNA origami we demonstrated that lbFCS+ yields truthful docking strand copy numbers *N* and dissociation/association rates *k*_off_ & *k*_on_ of the underlying imager/docking strand binding reaction from DNA-PAINT data sets acquired at moderate imager concentrations (≤ 5 nM) and measurement times ≤ 30 min). Our assessment of the working range indicated that lbFCS+ is suited for an application to localization clusters containing up to six docking strands. The high accuracy of lbFCS+ to determine hybridization rates allowed to measure small differences in imager binding dynamics to docking strands of same sequence but placed at different positions of nanoscopic DNA assemblies. Finally, this enabled us to resolve heterogeneous binding dynamics between individual DNA-PAINT clusters allowing for the distinction of stochastically generated and a priori indistinguishable DNA assemblies.

We want to stress that although lbFCS+ is in principle equally applicable to 3D data sets this work was limited to planar samples imaged in Total Internal Reflection Fluorescence (TIRF) configuration. Usually, a 3D DNA-PAINT image acquisition requires a confined illumination scheme (e.g. Highly Inclined and Laminated Optical Sheet (HILO) or Spinning Disc Confocal Microscopy (SDCM)) to suppress the fluorescent background from the imaging solution.

Since lbFCS+ is not relying on any ensemble averaging it would be ideally suited for the study of heterogeneous samples as expected in e.g. cellular environments. Heterogeneities might emerge from diffusional barriers due to compartmentalization or steric hindrance in densely-packed molecular environments. lbFCS+ could hence map the accessibility of imagers to different cellular parts (decoupled from the molecular copy numbers) which could be of general interest for the interpretation of DNA-PAINT images. Due to its accuracy in the determination of low molecular copy numbers especially studies aiming for the distinction of monomers, dimers or tetramers are a feasible first step. We reason that high target molecule densities[11], pronounced unspecific binding of imager strands [37, 45] and the optical sectioning capabilities of the used microscope will be major challenges when applying lbFCS+ to cellular targets. We want to highlight that DNA-PAINT (and hence lbFCS+) is generally not designed for living cells but requires fixed specimens. Finally, lbFCS+ requires the presence of well-separable localization clusters in the DNA-PAINT image and can as such not be readily transferred to an analysis of e.g. continuous objects.

We demonstrated that lbFCS+ is capable of detecting/distinguishing small differences in imager/docking strand binding dynamics in nanoscopic volumes containing low numbers of molecules requiring only moderate measurement times. Hence, lbFCS+ provides a highly-parallelized and easy-to-implement readout for potential on-chip bio-sensing applications. Especially interesting are applications requiring a direct detection of molecules in low concentration regimes without amplification steps [46, 47]. Additionally, our study of varying DNA-assemblies already suggests that lbFCS+ might readily serve as a readout to determine the state of logic gates (e.g. hairpins) in DNA-based logical circuits [48, 49]. However, we want to highlight that lbFCS+ is in principle not limited to the study of DNA hybridization reactions, but can applied to any reversible binding reaction of fluorescently-labeled ligands to immobilized receptors.

In conclusion, we believe that lbFCS+ provides a powerful tool with promising applications beyond its initial purpose of advancing quantitative DNA-PAINT imaging.

## AUTHOR CONTRIBUTIONS

J.S., F.S. and P.S. conceived the study. J.S. and F.S. designed and performed experiments. J.S. and F.S. designed and performed data analysis. F.S. wrote the analysis code. J.S., F.S. and P.S. wrote the manuscript. P.S. supervised the study. The co-first authorship order was determined via the best of three rounds in rock-paper-scissors at the start of the first authors’ PhD life. Both J.S and F.S. contributed equally. All authors discussed and interpreted results and revised the manuscript.

## ACKNOWLEDGMENTS

We thank Jan-Hagen Krohn for the implementation of the Chung-Kennedy filter in python and for design of the A20 adapter sequence. Further, we thank Sigrid Bauer and Sebastian Strauss for experimental support. J.S. and F.S. acknowledge support from Graduate School of Quantitative Bioscience Munich (QBM). All authors acknowledge support from the Center for Nano Science (CeNS). This work has been supported in part by the German Research Foundation through SFB1032 (projects A11 and A09 to R.J. and P.S.) and the Max Planck Society (R.J. and P.S.).

## SUPPLEMENTARY MATERIAL

Supplementary information can be found in a separate PDF on https://dummy.org

## REFERENCES

1. Stefan W Hell and Jan Wichmann. “Breaking the diffraction resolution limit by stimulated emission: stimulated-emission-depletion fluorescence microscopy”. In: Opt. Lett. 19.11 (1994), pp. 780–782. doi: 10.1364/OL.19.000780. url: http://ol.osa.org/abstract.cfm?URI=ol-19-11-780.

2. Thomas A Klar et al. “Fluorescence microscopy with diffraction resolution barrier broken by stimulated emission”. In: Proc. Natl. Acad. Sci. 97.15 (2000), 8206 LP–8210. doi: 10.1073/pnas.97.15.8206. url: http://www.pnas.org/content/97/15/8206.abstract.

3. Eric Betzig et al. “Imaging Intracellular Fluorescent Proteins at Nanometer Resolution”. In: Science (80-.). 313.5793 (2006), 1642 LP–1645. doi: 10.1126/science.1127344. url: http://science.sciencemag.org/content/313/5793/1642.abstract.

4. Michael J Rust, Mark Bates, and Xiaowei Zhuang. “Sub-diffraction-limit imaging by stochastic optical reconstruction microscopy (STORM)”. In: Nat. Methods 3.10 (2006), pp. 793–796. issn: 1548-7105. doi: 10.1038/nmeth929. url: https://doi.org/10.1038/nmeth929.

5. Alexey Sharonov and Robin M Hochstrasser. “Wide-field subdiffraction imaging by accumulated binding of diffusing probes”. In: Proc. Natl. Acad. Sci. 103.50 (2006), 18911 LP–18916. doi: 10.1073/pnas.0609643104. url: http://www.pnas.org/content/103/50/18911.abstract.

6. Ralf Jungmann et al. “Single-Molecule Kinetics and Super-Resolution Microscopy by Fluorescence Imaging of Transient Binding on DNA Origami”. In: Nano Lett. 10.11 (2010), pp. 4756–4761. issn: 1530-6984. doi: 10.1021/nl103427w. url: https://doi.org/10.1021/nl103427w.

7. Joerg Schnitzbauer et al. “Super-resolution microscopy with DNA-PAINT”. In: Nat. Protoc. 12.6 (2017), pp. 1198–1228. issn: 1750-2799. doi: 10.1038/nprot.2017.024. url: https://doi.org/10.1038/nprot.2017.024.

8. Russell E Thompson, Daniel R Larson, and Watt W Webb. “Precise Nanometer Localization Analysis for Individual Fluorescent Probes”. In: Biophys. J. 82.5 (2002), pp. 2775–2783. issn: 0006-3495. doi: https://doi.org/10.1016/S0006-3495(02)75618-X. url: https://www.sciencedirect.com/science/article/pii/S000634950275618X.

9. Hendrik Deschout et al. “Precisely and accurately localizing single emitters in fluorescence microscopy”. In: Nat. Methods 11.3 (2014), pp. 253–266. issn: 1548-7105. doi: 10.1038/nmeth.2843. url: https://doi.org/10.1038/nmeth.2843.

10. Sebastian Strauss and Ralf Jungmann. “Up to 100-fold speed-up and multiplexing in optimized DNA-PAINT”. In: Nat. Methods 17.8 (2020), pp. 789–791. issn: 1548-7105. doi: 10.1038/s41592-020-0869-x. url: https://doi.org/10.1038/s41592-020-0869-x.

11. David Baddeley and Joerg Bewersdorf. “Biological Insight from Super-Resolution Microscopy: What We Can Learn from Localization-Based Images”. In: Annu. Rev. Biochem. 87.1 (2018), pp. 965–989. issn: 0066-4154. doi: 10.1146/annurev-biochem-060815-014801. url: https://doi.org/10.1146/annurev-biochem-060815-014801.

12. Paolo Annibale et al. “Quantitative Photo Activated Localization Microscopy: Unraveling the Effects of Photoblinking”. In: PLoS One 6.7 (2011), e22678. url: https://doi.org/10.1371/journal.pone.0022678.

13. Paolo Annibale et al. “Identification of clustering artifacts in photoactivated localization microscopy”. In: Nat. Methods 8.7 (2011), pp. 527–528. issn: 1548-7105. doi: 10.1038/nmeth.1627. url: https://doi.org/10.1038/nmeth.1627.

14. Carla Coltharp, Rene P Kessler, and Jie Xiao. “Accurate Construction of Photoactivated Localization Microscopy (PALM) Images for Quantitative Measurements”. In: PLoS One 7.12 (2012), e51725. url: https://doi.org/10.1371/journal.pone.0051725.

15. Sang-Hyuk Lee et al. “Counting single photoactivatable fluorescent molecules by photoactivated localization microscopy (PALM)”. In: Proc. Natl. Acad. Sci. 109.43 (2012), 17436 LP–17441. doi: 10.1073/pnas.1215175109. url: http://www.pnas.org/content/109/43/17436.abstract.

16. Elias M Puchner et al. “Counting molecules in single organelles with superresolution microscopy allows tracking of the endosome maturation trajectory”. In: Proc. Natl. Acad. Sci. 110.40 (2013), 16015 LP–16020. doi: 10.1073/pnas.1309676110. url: http://www.pnas.org/content/110/40/16015.abstract.

17. Xiaolin Nan et al. “Single-molecule superresolution imaging allows quantitative analysis of RAF multimer formation and signaling”. In: Proc. Natl. Acad. Sci. 110.46 (2013), 18519 LP–18524. doi: 10.1073/pnas.1318188110. url: http://www.pnas.org/content/110/46/18519.abstract.

18. Nadine Ehmann et al. “Quantitative super-resolution imaging of Bruchpilot distinguishes active zone states”. In: Nat. Commun. 5.1 (2014), p. 4650. issn: 2041-1723. doi: 10.1038/ncomms5650. url: https://doi.org/10.1038/ncomms5650.

19. Geoffrey C Rollins et al. “Stochastic approach to the molecular counting problem in superresolution microscopy”. In: Proc. Natl. Acad. Sci. 112.2 (2015), E110 LP–E118. doi: 10.1073/pnas.1408071112. url: http://www.pnas.org/content/112/2/E110.abstract.

20. Franziska Fricke et al. “One, two or three? Probing the stoichiometry of membrane proteins by single-molecule localization microscopy”. In: Sci. Rep. 5.1 (2015), p. 14072. issn: 2045-2322. doi: 10.1038/srep14072. url: https://doi.org/10.1038/srep14072.

21. Maria Aurelia Ricci et al. “Chromatin Fibers Are Formed by Heterogeneous Groups of Nucleosomes In Vivo”. In: Cell 160.6 (2015), pp. 1145–1158. issn: 0092-8674. doi: https://doi.org/10.1016/j.cell.2015.01.054. url: https://www.sciencedirect.com/science/article/pii/S0092867415001324.

22. Gerhard Hummer, Franziska Fricke, and Mike Heilemann. “Model-independent counting of molecules in single-molecule localization microscopy”. In: Mol. Biol. Cell 27.22 (2016), pp. 3637–3644. issn: 1059-1524. doi: 10.1091/mbc.E16-07-0525. url: https://doi.org/10.1091/mbc.E16-07-0525.

23. Caroline Laplante et al. “Molecular organization of cytokinesis nodes and contractile rings by super-resolution fluorescence microscopy of live fission yeast”. In: Proc. Natl. Acad. Sci. 113.40 (2016), E5876 LP–E5885. doi: 10.1073/pnas.1608252113. url: http://www.pnas.org/content/113/40/E5876.abstract.

24. Daniel Nino et al. “Molecular Counting with Localization Microscopy: A Bayesian Estimate Based on Fluorophore Statistics”. In: Biophys. J. 112.9 (2017), pp. 1777–1785. issn: 0006-3495. doi: https://doi.org/10.1016/j.bpj.2017.03.020. url: https://www.sciencedirect.com/science/article/pii/S0006349517303399.

25. Philip R Nicovich, Dylan M Owen, and Katharina Gaus. “Turning single-molecule localization microscopy into a quantitative bioanalytical tool”. In: Nat. Protoc. 12.3 (2017), pp. 453–460. issn: 1750-2799. doi: 10.1038/nprot.2016.166. url: https://doi.org/10.1038/nprot.2016.166.

26. Ottavia Golfetto et al. “A Platform To Enhance Quantitative Single Molecule Localization Microscopy”. In: J. Am. Chem. Soc. 140.40 (2018), pp. 12785–12797. issn: 0002-7863. doi: 10.1021/jacs.8b04939. url: https://doi.org/10.1021/jacs.8b04939.

27. Samuel T Hess, Thanu P K Girirajan, and Michael D Mason. “Ultra-High Resolution Imaging by Fluorescence Photoactivation Localization Microscopy”. In: Biophys. J. 91.11 (2006), pp. 4258–4272. issn: 0006-3495. doi: https://doi.org/10.1529/biophysj.106.091116. url: https://www.sciencedirect.com/science/article/pii/S0006349506721403.

28. Mike Heilemann et al. “Subdiffraction-Resolution Fluorescence Imaging with Conventional Fluorescent Probes”. In: Angew. Chemie Int. Ed. 47.33 (2008), pp. 6172–6176. issn: 1433-7851. doi: https://doi.org/10.1002/anie.200802376. url: https://doi.org/10.1002/anie.200802376.

29. Alexander P Demchenko. “Photobleaching of organic fluorophores: quantitative characterization, mechanisms, protection”. In: Methods Appl. Fluoresc. 8.2 (2020), p. 22001. issn: 2050-6120. doi: 10.1088/2050-6120/ab7365. url: http://dx.doi.org/10.1088/2050-6120/ab7365.

30. Philipp Blumhardt et al. Photo-Induced Depletion of Binding Sites in DNA-PAINT Microscopy. 2018. doi: 10.3390/molecules23123165.

31. F. Stehr et al. “Flat-top TIRF illumination boosts DNA-PAINT imaging and quantification”. In: Nat. Commun. 10.1 (2019). issn: 20411723. doi: 10.1038/s41467-019-09064-6.

32. J. Stein et al. “Toward Absolute Molecular Numbers in DNA-PAINT”. In: Nano Lett. 19.11 (2019). issn: 15306992. doi: 10.1021/acs.nanolett.9b03546.

33. Mickaël Lelek et al. “Single-molecule localization microscopy”. In: Nat. Rev. Methods Prim. 1.1 (2021), p. 39. issn: 2662-8449. doi: 10.1038/s43586-021-00038-x. url: https://doi.org/10.1038/s43586-021-00038-x.

34. Ralf Jungmann et al. “Quantitative super-resolution imaging with qPAINT”. In: Nat. Methods 13.5 (2016), pp. 439–442. issn: 1548-7105. doi: 10.1038/nmeth.3804. url: https://doi.org/10.1038/nmeth.3804.

35. Matthias Schickinger, Martin Zacharias, and Hendrik Dietz. “Tethered multifluorophore motion reveals equilibrium transition kinetics of single DNA double helices”. In: Proc. Natl. Acad. Sci. 115.32 (2018), E7512 LP–E7521. doi: 10.1073/pnas.1800585115. url: http://www.pnas.org/content/115/32/E7512.abstract.

36. F. Schueder et al. “An order of magnitude faster DNA-PAINT imaging by optimized sequence design and buffer conditions”. In: Nat. Methods 16.11 (2019). issn: 15487105. doi: 10.1038/s41592-019-0584-7.

37. Alexander H Clowsley et al. “Repeat DNA-PAINT suppresses background and non-specific signals in optical nanoscopy”. In: Nat. Commun. 12.1 (2021), p. 501. issn: 2041-1723. doi: 10.1038/s41467-020-20686-z. url: https://doi.org/10.1038/s41467-020-20686-z.

38. Paul W K Rothemund. “Folding DNA to create nanoscale shapes and patterns”. In: Nature 440.7082 (2006), pp. 297–302. issn: 1476-4687. doi: 10.1038/nature04586. url: https://doi.org/10.1038/nature04586.

39. Florian Stehr et al. “Tracking single particles for hours via continuous DNA-mediated fluorophore exchange”. In: Nat. Commun. 12.1 (2021), p. 4432. issn: 2041-1723. doi: 10.1038/s41467-021-24223-4. url: https://doi.org/10.1038/s41467-021-24223-4.

40. S H Chung and R A Kennedy. “Forward-backward non-linear filtering technique for extracting small biological signals from noise”. In: J. Neurosci. Methods 40.1 (1991), pp. 71–86. issn: 0165-0270. doi: https://doi.org/10.1016/0165-0270(91)90118-J. url: https://www.sciencedirect.com/science/article/pii/016502709190118J.

41. Jonas Mücksch et al. “Quantifying Reversible Surface Binding via Surface-Integrated Fluorescence Correlation Spectroscopy”. In: Nano Lett. 18.5 (2018), pp. 3185–3192. issn: 1530-6984. doi: 10.1021/acs.nanolett.8b00875. url: https://doi.org/10.1021/acs.nanolett.8b00875.

42. Arthur D Edelstein et al. “Advanced methods of microscope control using *μ*Manager software”. In: J. Biol. Methods 1.2 SE - Protocols (2014), e10. doi: 10.14440/jbm.2014.36. url: https://jbmethods.org/jbm/article/view/36.

43. Sebastian Strauss et al. “Modified aptamers enable quantitative sub-10-nm cellular DNA-PAINT imaging”. In: Nat. Methods 15.9 (2018), pp. 685–688. issn: 1548-7105. doi: 10.1038/s41592-018-0105-0. url: https://doi.org/10.1038/s41592-018-0105-0.

44. Leland McInnes, John Healy, and Steve Astels. “hdbscan: Hierarchical density based clustering”. In: J. Open Source Softw. 2.11 (2017),p. 205. doi: 10.21105/joss.00205. url: https://doi.org/10.21105/joss.00205.

45. H J Geertsema et al. “Left-handed DNA-PAINT for improved super-resolution imaging in the nucleus”. In: Nat. Biotechnol. 39.5 (2021), pp. 551–554. issn: 1546-1696. doi: 10.1038/s41587-020-00753-y. url: https://doi.org/10.1038/s41587-020-00753-y.

46. Alexander Johnson-Buck et al. “A guide to nucleic acid detection by single-molecule kinetic fingerprinting”. In: Methods 153 (2019), pp. 3–12. issn: 1046-2023. doi: https://doi.org/10.1016/j.ymeth.2018.08.002. url: https://www.sciencedirect.com/science/article/pii/S1046202318300951.

47. Tanmay Chatterjee et al. “Direct kinetic fingerprinting and digital counting of single protein molecules”. In: Proc. Natl. Acad. Sci. 117.37 (2020), 22815 LP–22822. doi: 10.1073/pnas.2008312117. url: http://www.pnas.org/content/117/37/22815.abstract.

48. Lulu Qian and Erik Winfree. “Scaling Up Digital Circuit Computation with DNA Strand Displacement Cascades”. In: Science (80-.). 332.6034 (2011), 1196 LP–1201. doi: 10.1126/science.1200520. url: http://science.sciencemag.org/content/332/6034/1196.abstract.

49. Kevin M Cherry and Lulu Qian. “Scaling up molecular pattern recognition with DNA-based winner-take-all neural networks”. In: Nature 559.7714 (2018), pp. 370–376. issn: 1476-4687. doi: 10.1038/s41586-018-0289-6. url: https://doi.org/10.1038/s41586-018-0289-6.

