## Supplementary Information for "Calibration free counting of low molecular copy numbers in single DNA-PAINT localization clusters"

#### SUPPLEMENTARY FIGURES

1 **Intensity denoising and normalization.** . . . . . 2
2 **Exemplary intensity traces ($N = 4$) corresponding to selected intervals in $\hat{I}$.** . . . . . 3
3 **Calibration of $k_{on}$ using Pm2* reference origami.** . . . . . 4
4 **lbFCS+ results for $N = 12$ origami measured at an imager concentration of 2.5 nM.** . . . . . 4
5 **Control for position dependency of $k_{off}$&$k_{on}$.** . . . . . 5
6 **Selection of intervals in $\hat{I}$ for $N = 4$ origami measured at imager concentration of 1.25 nM.** . . . . . 5
7 **Computational combination of $N = 1$ clusters measured at imager concentration of 1.25 nM.** . . . . . 6
8 **Quantitative imaging of 18 FOVs in 3 h.** . . . . . 6
9 **2xLink origami incubated with varying linker strand concentrations.** . . . . . 7

#### SUPPLEMENTARY TABLES

1 **Detailed imaging conditions.** . . . . . 8

### SUPPLEMENTARY FIGURES

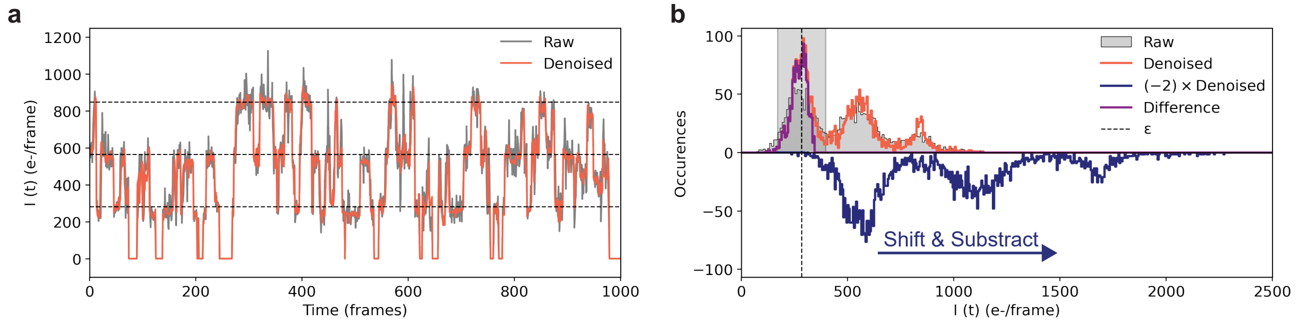


**Supplementary Figure 1: Intensity denoising and normalization.** **(a)** Exemplary raw intensity trace from a localization cluster from a data set containing $N = 4$ DNA origami(grey). The intensity is given by the total number of photo-electrons contained in the fitted 2D Gaussian PSF during each single molecule localization event, i.e., (# photo-electrons)/frame. The intensity trace exhibits step-like behavior between three intensity levels due to varying numbers of imager strands that are bound to the origami at a given time (i.e., per frame). Smaller intensity fluctuations at each level are due to the noise induced by the camera electronics (readout noise) and due to the stochastic nature of the photon arrivals (shot noise). To regain a close to step-like behavior according to the number of simultaneously bound imagers, a non-linear de-noising filter [1] is applied to the raw intensity trace during analysis; the de-noised intensity trace is shown in red. The dashed lines correspond to multiples of the normalization value $\epsilon$ as obtained by the following procedure. **(b)** Photo-electron intensity histograms as obtained from the raw intensity trace (grey) and de-noised intensity trace (red) as shown in (a) both feature three clearly distinguishable peaks, indicating that up to three imagers are bound simultaneously to the origami. The normalization process is based on the histogram of the de-noised intensity trace. Its general idea is to isolate the leftmost peak of the histogram in order to obtain the (cluster specific) intensity value corresponding to a single imager bound to the target. Therefore, the intensity values of the de-noised intensity trace are first multiplied by a a factor of $\times 2$ and the histogram is re-calculated maintaining the original bin size of the de-noised histogram. Subsequently the resulting bin heights are adjusted to approximately match the heights of the original histogram by multiplication with a factor of $\times 1.5$. Finally, multiplication of the bin heights with a factor of $\times(-1)$ yields the '$(-2)\times$De-noised' histogram as shown in blue. Now, the '$(-2)\times$De-noised' histogram is subtracted from the original histogram (de-noised; red) and subsequently shifted one bin to the right. This 'Shift & Substract' procedure is repeated $\times 100$ yielding the difference histogram (turquoise) only consisting of the original leftmost peak. As a next step, all values in the original de-noised histogram not deviating more than 40 % to the median of the difference histogram are selected (light-grey interval). Finally, the intensity value $\epsilon$ (dashed line) used for normalization is set to the median of the de-noised intensity values within the selected interval.

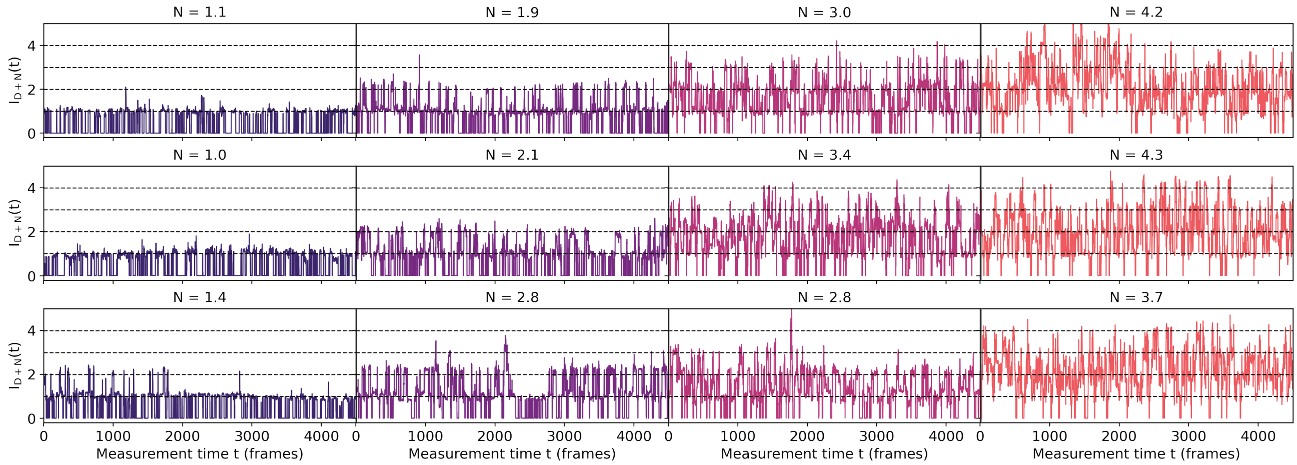


**Supplementary Figure 2: Exemplary intensity traces ($N = 4$) corresponding to selected intervals in $\hat{I}$.** Exemplary de-noised and normalized intensity traces as obtained from $N = 4$ origami according to the selected intervals in $\hat{I}$ as depicted in the left panel of **Fig.2e** (i.e., same color code used for traces and intervals in $\hat{I}$). The counting result $N$ as obtained by lbFCS+ analysis is stated above each trace. As expected, observed intensity levels are located at unit-less integer values (i.e.,: 1, 2, 3, 4) after the normalization procedure (described in **Supplementary Fig.1** according to the current number of imagers bound to the DNA origami (dashed lines).

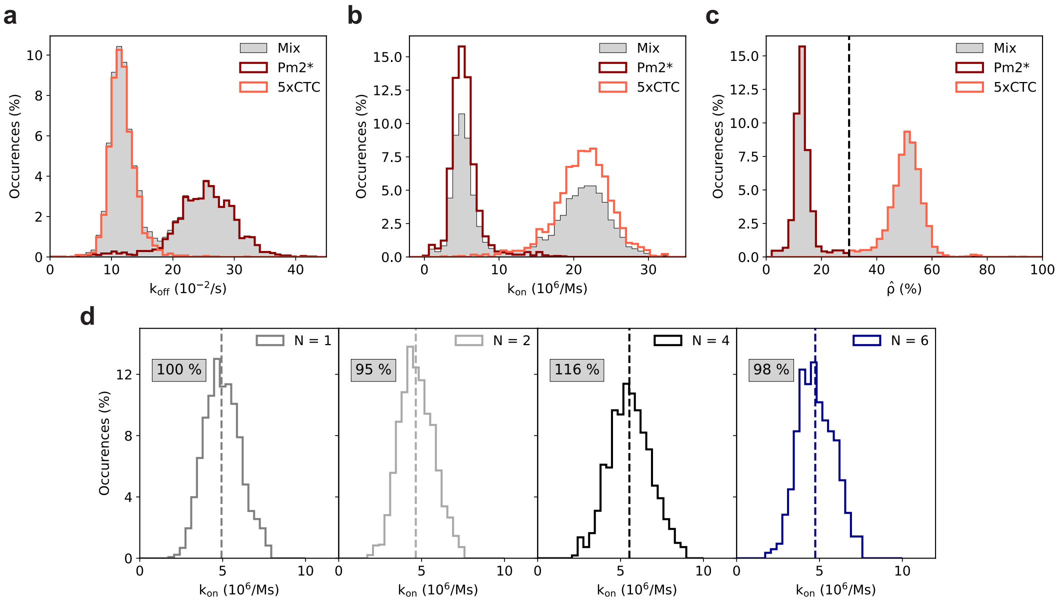


**Supplementary Figure 3: Calibration of $k_{on}$ using Pm2* reference origami. (a)** Resulting $k_{off}$ distribution from a sample containing a mixture of origami with either a single (i.e., $N = 1$) 5xCTC docking strand or a single Pm*2 docking strand (calibration reference) imaged with the same imager Pm2 at a concentration of 5 nM (grey; 'Mix'). Since imagers bind to Pm2* docking strands with a higher $k_{off}$ and lower $k_{on}$ (due to a repetitive docking strand design, described in refs. [2–5]) the individual distributions corresponding to either Pm2*(dark red) or 5xCTC origami (orange) can be easily distinguished using the occupancy $\hat{\rho}$; see (c). The mixed sample contained a total of # clusters = 4,076 with # clusters = 1,833 assigned to Pm2*, and # clusters = 2,243 assigned to 5xCTC. **(b)** Same data and analogous analysis as in (a) but for the association rate $k_{on}$. **(c)** Illustration of the isolation of clusters based on the occupancy $\hat{\rho}$ (dashed line), i.e., $\hat{\rho} < 30$ % corresponds to Pm2* origami (dark red) and $\hat{\rho} > 30$ % corresponds to 5xCTC origami (orange); as used in (a,b). **(d)** As described in Section 2.4 each of the samples shown in **Fig.1a-d,f** contained a subpopulation of Pm2* reference origami in order to track pipetting errors of the imager concentration affecting the resulting $k_{on} = \tilde{k}_{on}/c$. We first isolated the Pm2* clusters in each sample according to the procedure described in (a-c). Subsequently we compared the resulting median $k_{on}$ (dashed line) of each subpopulation (same color code as in **Fig.1a-d,f**) to a fixed value of $\overline{k_{on}} \equiv 5 \times 10^6$ 1/Ms. In the final calibration step, the relative deviation of the median $k_{on}$ for each Pm2* subpopulation to $\overline{k_{on}}$ (grey boxes) was then used to correct the $k_{on}$ as obtained from the target 5xCTC $N = 1, 2, 4, 6$ origami. Pm2* reference origami subpopulations contained: # clusters = 1,330 for $N = 1$, # clusters = 1,920 for $N = 2$, # clusters = 1,098 for $N = 4$, # clusters = 845 for $N = 6$.

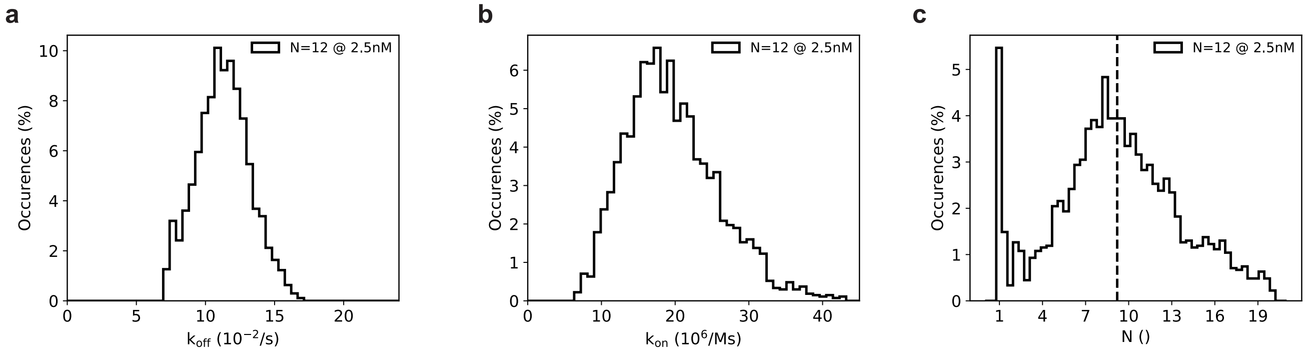


**Supplementary Figure 4: lbFCS+ results for $N = 12$ origami measured at an imager concentration of 2.5 nM. (a)** lbFCS+ results for $k_{off}$. **(b)** lbFCS+ results for $k_{on}$. **(c)** lbFCS+ results for $N$. Based on a mean of $\langle N \rangle = 9.2$ (dashed line) we estimated an average incorporation efficiency of 77 % (in good agreement with [6]). Data shown in (a,b,c) contained a total of # clusters = 2,689.

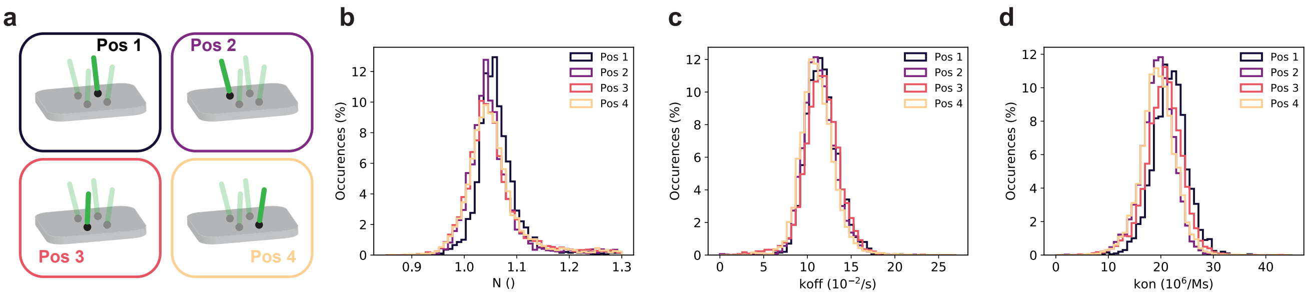


**Supplementary Figure 5: Control for position dependency of $k_{\mathrm{off}}$&$k_{\mathrm{on}}$. (a)** To check for position dependent hybridization rates we folded DNA origami each containing only one of the four docking strands (i.e., $N$ = 1) at every position used for the $N$ = 4 origami (Pos1-4; transparent docking strands indicate other $N$ = 4 positions). **(b)** Corresponding lbFCS+ results for $N$ for all origami shown in (a) agree with the expected $N$ = 1. **(c)** Corresponding lbFCS+ results for $k_{\mathrm{off}}$ for all origami shown in (a) did not indicate any position dependent alteration of $k_{\mathrm{off}}$. **(d)** Corresponding lbFCS+ results for $k_{\mathrm{on}}$ for all origami shown in (a) did not indicate any position dependent alteration of $k_{\mathrm{on}}$. Data shown in (b,c,d) contained # clusters = 1,994 for Pos1, # clusters = 3,188 for Pos2, # clusters = 6,428 for Pos3, # clusters = 5,680 for Pos4.

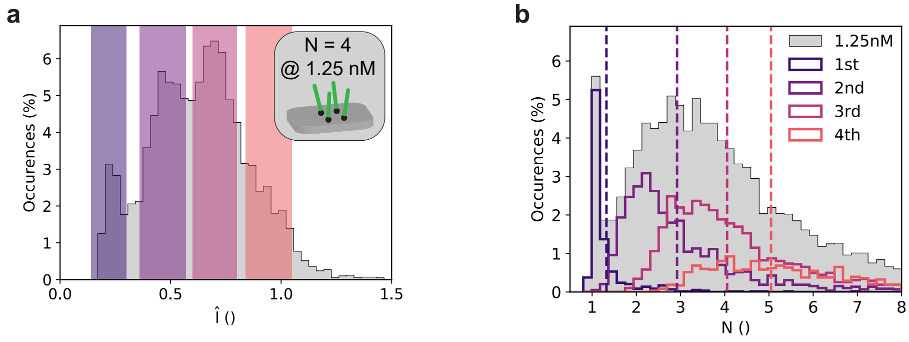


**Supplementary Figure 6: Selection of intervals in $\hat{I}$ for $N$ = 4 origami measured at imager concentration of 1.25 nM. (a)** Selection of intervals in $\hat{I}$, analogous to the left panels of **Fig.2e** and **Fig.3c**. **(b)** Inspection of the counting results for clusters corresponding to the selected intervals in $\hat{I}$ as shown in (a), analogous to the right panels of **Fig.2e** and **Fig.3c**. Data shown in (a,b) contained # clusters = 5,834.

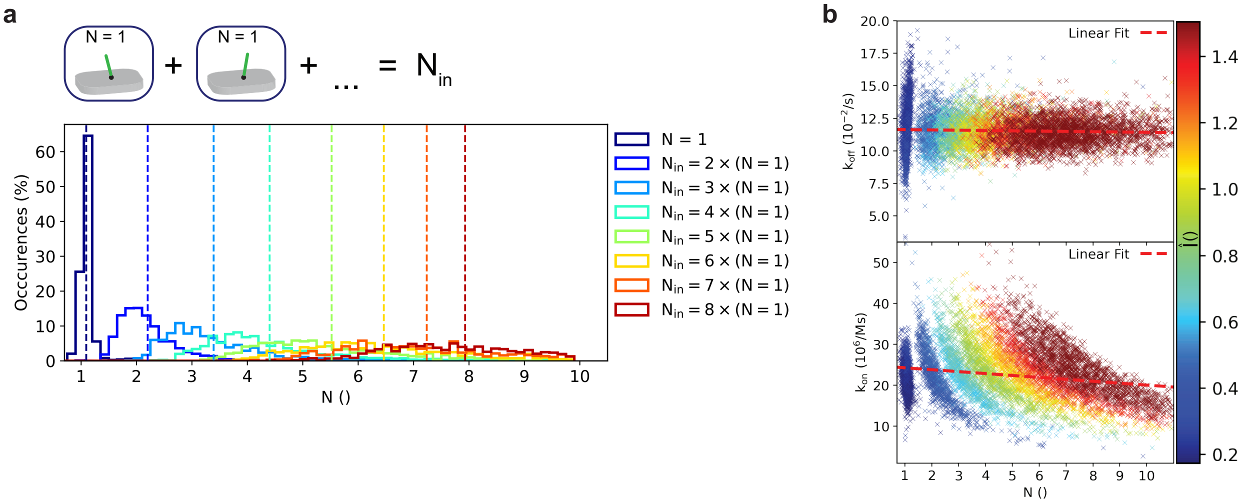


**Supplementary Figure 7: Computational combination of $N = 1$ clusters measured at imager concentration of 1.25 nM.** **(a)** lbFCS+ counting results obtained from computationally regrouped clusters consisting of up to eight experimental $N = 1$ clusters measured at an imager concentration of 1.25 nM (i.e., $N_{\text{in}} = k \times (N = 1)$ up to $k = 8$; # clusters = 1,000 for each $N_{\text{in}}$), analogous to **Fig.3a**. **(b)** $k_{\text{off}}$ & $k_{\text{on}}$ vs. $N$ scatter plot (color coded by $\hat{I}$) and linear fit (red dashed line) of the same data as shown in (a), analogous to the left panels of **Fig.3b**. Data shown in (b) contained a total of # clusters = 8,648.

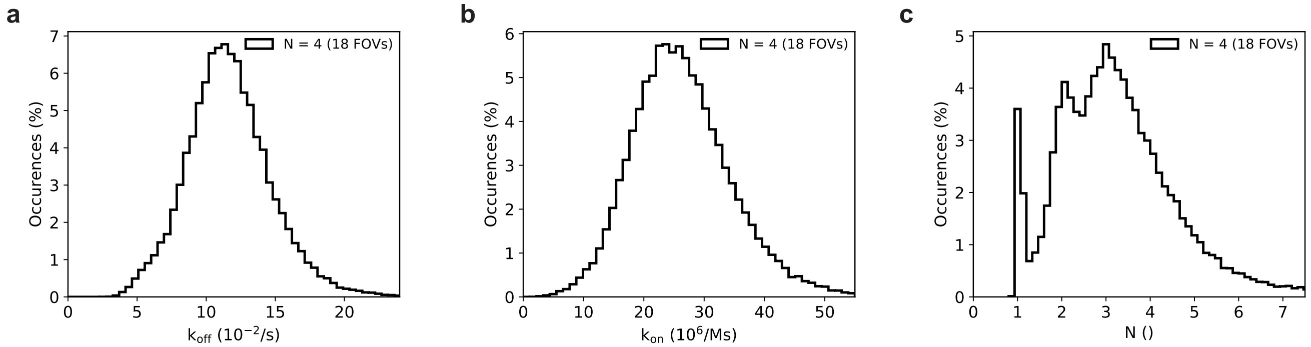


**Supplementary Figure 8: Quantitative imaging of 18 FOVs in 3 h.** **(a)** lbFCS+ results for $k_{\text{off}}$ as obtained from a total of 18 FOVs of a sample containing $N = 4$ origami (total # clusters = 48,966). Each FOV was measured for 1500 frames (i.e., 10 min) leading to a total of 3 h of measurement time for all 18 FOVs. **(b)** Same as (a) but for $k_{\text{on}}$. We want to highlight that this sample did not contain a subpopulation of Pm2* origami used for calibration of $k_{\text{on}}$. **(c)** Same as (a) but for $N$.

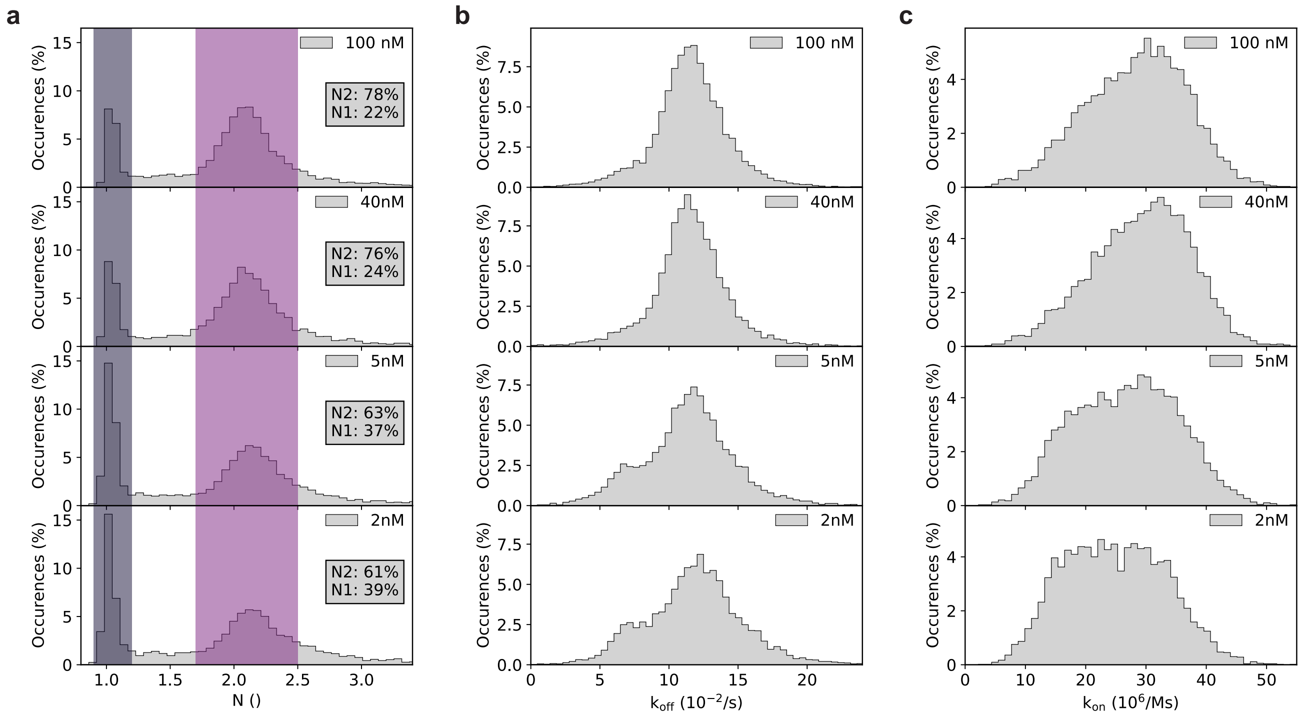


**Supplementary Figure 9: 2xLink origami incubated with varying linker strand concentrations. (a)** lbFCS+ counting results for 2xLink origami incubated for 3 min at linker strand concentrations c = 100 nM, 40 nM, 5 nM, 2 nM prior to imaging. We selected 2xLink origami yielding $0.9 < N < 1.2$ corresponding to the $N = 1$ configurations (left colored interval) as shown in the blue box of **Fig.4e**, and 2xLink origami yielding $1.7 < N < 2.5$ corresponding to the $N = 2$ configuration (right colored interval) as shown in the dark-red box of **Fig.4e**. The grey boxes indicate the ratio of 2xLink origami found in either a $N = 1$ or $N = 2$ state (normalized to the total of origami in a $N = 1$ or $N = 2$ state). **(b)** lbFCS+ result for $k_{\mathrm{off}}$ for the same data as shown in (a). **(c)** lbFCS+ result for $k_{\mathrm{on}}$ for the same data as shown in (a). Data shown in (a,b,c) contained # clusters = 9,372 for 100 nM, # clusters = 5,625 for 40 nM, # clusters = 6,709 for 5 nM, # clusters = 5,298 for 2 nM.

#### SUPPLEMENTARY TABLES

**Supplementary Table 1: Detailed imaging conditions.** For all presented data the exposure time was set to 400 ms corresponding to the image aquisition duty cycle (see Section 2.5). ‘Reference Pm2*’ indicates if the sample contained - besides the target 5xCTC origami - a Pm2* ($N = 1$) origami subpopulation for calibration of $k_{on}$ (see Section 2.4). In case of ‘Link’ and ‘2xLink’ origami the given concentration indicates the concentration of linker strands during the the 3 min incubation time prior to imaging.

| Figure | Sample | Reference Pm2* | Imager concentration (nM) | Irradiance (W/cm$^2$) | Frames |
|---|---|---|---|---|---|
| Fig.2<br>Fig.3b (right panel)<br>SI_Fig.3 | N = 1,2,4,6 | Yes | 5 | 10 | 4500 |
| SI_Fig.1<br>SI_Fig.2<br>Fig.3d,e | N = 4 | Yes | 5 | 10 | 4500 |
| Fig.3a<br>Fig.3b (left panel) | N = 1, regrouped | Yes | 5 | 10 | 4500 |
| SI_Fig.4 | N = 12 | Yes | 2.5 | 10 | 4500 |
| SI_Fig.5 | N = 1, Pos1-4 | Yes | 5 | 10 | 4500 |
| Fig.3c<br>Fig.3d,e | N = 4 | Yes | 2.5 | 10 | 4500 |
| SI_Fig.6<br>Fig.3d,e | N = 4 | Yes | 1.25 | 10 | 4500 |
| SI_Fig.7 | N = 1, regrouped | No | 1.25 | 10 | 4500 |
| Fig.3f,g,h | N=4 | Yes | 5 | 10 | 4500<br>2250<br>1125<br>600 |
| SI_Fig.8 | N = 4, 18 FOVs | No | 5 | 10 | 1500 |
| Fig.4b,c,d | N=1, Direct<br>Link, 100 nM<br>Direct + Link | Yes | 10<br>10<br>5 | 10 | 4500 |
| Fig.4f,g,h,i<br>SI_Fig.9 | 2xLink, 100 nM<br>2xLink, 40 nM<br>2xLink, 5 nM<br>2xLink, 2 nM | Yes | 5 | 10 | 4500 |

#### REFERENCES


1. S H Chung and R A Kennedy. “Forward-backward non-linear filtering technique for extracting small biological signals from noise”. In: *J. Neurosci. Methods* 40.1 (1991), pp. 71–86. ISSN: 0165-0270. DOI: https://doi.org/10.1016/0165-0270(91)90118-J. URL: https://www.sciencedirect.com/science/article/pii/016502709190118J.
2. J. Stein et al. “Toward Absolute Molecular Numbers in DNA-PAINT”. In: *Nano Lett.* 19.11 (2019). ISSN: 15306992. DOI: 10.1021/acs.nanolett.9b03546.
3. Sebastian Strauss and Ralf Jungmann. “Up to 100-fold speed-up and multiplexing in optimized DNA-PAINT”. In: *Nat. Methods* 17.8 (2020), pp. 789–791. ISSN: 1548-7105. DOI: 10.1038/s41592-020-0869-x. URL: https://doi.org/10.1038/s41592-020-0869-x.
4. Alexander H Clowsley et al. “Repeat DNA-PAINT suppresses background and non-specific signals in optical nanoscopy”. In: *Nat. Commun.* 12.1 (2021), p. 501. ISSN: 2041-1723. DOI: 10.1038/s41467-020-20686-z. URL: https://doi.org/10.1038/s41467-020-20686-z.
5. Florian Stehr et al. “Tracking single particles for hours via continuous DNA-mediated fluorophore exchange”. In: *Nat. Commun.* 12.1 (2021), p. 4432. ISSN: 2041-1723. DOI: 10.1038/s41467-021-24223-4. URL: https://doi.org/10.1038/s41467-021-24223-4.

6. Sebastian Strauss et al. "Modified aptamers enable quantitative sub-10-nm cellular DNA-PAINT imaging". In: *Nat. Methods* 15.9 (2018), pp. 685–688. ISSN: 1548-7105. DOI: 10.1038/s41592-018-0105-0. URL: https://doi.org/10.1038/s41592-018-0105-0.
